# A variable-gain stochastic pooling motif mediates information transfer from receptor assemblies into NF-κB

**DOI:** 10.1101/2021.03.29.437543

**Authors:** J. Agustin Cruz, Chaitanya S. Mokashi, Gabriel J. Kowalczyk, Yue Guo, Qiuhong Zhang, Sanjana Gupta, David L. Schipper, Robin E. C. Lee

**Affiliations:** Department of Computational and Systems Biology, School of Medicine, University of Pittsburgh, Pittsburgh, PA 15213, USA; Department of Physics and Astronomy, University of Pittsburgh, Pittsburgh, PA 15260, USA

## Abstract

A myriad of inflammatory cytokines regulate signaling pathways to maintain cellular homeostasis. The IKK complex is an integration hub for cytokines that govern NF-κB signaling. In response to inflammation, IKK is activated through recruitment to receptor-associated protein assemblies. How and what information IKK complexes transmit about the milieu are open questions. Here we track dynamics of IKK complexes and nuclear NF-κB to identify upstream signaling features that determine same-cell responses. Experiments and modeling of single complexes reveals their size, number, and timing relays cytokine-specific information with feedback control that is independent of transcription. Our results provide evidence for variable-gain stochastic pooling, a noise-reducing motif that enables parsimonious and cytokine-specific information transfer. We propose that emergent properties of stochastic pooling are general principles of receptor signaling.

**One Sentence Summary:** A variable-gain stochastic pooling motif mediates robust and tunable information transmission from the extracellular milieu into the cell.

## Main Text

### Introduction

A limited number of transmembrane receptors expressed on the cell surface mediate crucial transmission of information between extracellular and intracellular signaling molecules. Key questions are understanding the mechanisms and limitations that underlie signal transmission, in particular for cytokine receptor signaling that is often deregulated in disease. The nuclear-factor kappa-B (NF-κB) signaling pathway is an archetypal molecular communication channel that transmits information about extracellular cytokines to regulate cellular adaptation through activation of the RelA transcription factor (*1-3*). When ligated with inflammatory molecules, such as tumor necrosis factor (TNF), interleukin-1-β (IL-1), among others, many activated receptors converge on NF-κB signaling (*4, 5*).

Ligation of TNF to the TNF receptor (TNFR1) recruits adaptor proteins and enzymes to form a large multiprotein complex near the plasma membrane (*6-9*). Ubiquitin modifying enzymes are critical components that assemble linear, branched, and mixed polyubiquitin scaffolds around the multiprotein complex (*10-13*). The NEMO subunit of the cytoplasmic IκB-kinase (IKK) complex is rapidly recruited via direct interaction with the polyubiquitin scaffold and accessory proteins, where IKK is activated through induced proximity with regulatory kinases (*4, 14-16*). The fully assembled TNFR1 complex, referred to as ‘Complex I’ (CI; (*6*)), is a master regulator of inflammation-dependent NF-κB signaling. Although other inflammatory molecules such as IL-1 signal through CI-like complexes using different receptors, adaptor proteins (*17, 18*), and varying compositions of ubiquitin chain scaffolds (*19, 20*), all rely on induced-proximity activation of IKK in regulation of NF-κB (*21*).

When observed in single cells exposed to inflammatory stimuli, the RelA subunit of NF-κB encodes a dynamic transcriptional signal by translocating from the cytoplasm into the nucleus (*2, 3, 22-24*). Models calibrated to single-cell RelA data (*23-26*) have revealed numerous transcriptional mechanisms and emergent properties that place the NF-κB pathway among exemplars of dynamical biological systems (*27, 28*). Key to these findings are two mediators of negative feedback, IκBα and A20, which are transcriptionally regulated by NF-κB. IκBα restores NF-κB to its baseline cytoplasmic localization through nuclear export and sequestration, whereas A20 diminishes kinase activation upstream of NF-κB through disassembly of CI-like structures in addition to non-catalytic mechanisms (*10, 23, 25, 26, 29*). Dynamical regulation of transcription and feedback via NF-κB is strongly recapitulated between models and experiments; however, there is a dearth of quantitative single-cell data at the level of cytokine detection and dynamical properties of CI-like complexes to substantiate our understanding of upstream signal transmission.

Here, we develop genetically modified cells that endogenously express fluorescent protein fusions of NEMO and RelA, allowing same-cell measurements of CI-like structures and canonical NF-kB signaling from live-cell images. We establish differences between TNF and IL1 responses in biophysical properties of NEMO complexes and demonstrate a continuum relating CI-like structures and downstream NF-kB responses in the same cell. By tracking single complexes, we demonstrate that: i) cytokine dosage and time-varying presentation modulates the timing and numbers of CI-like structures; ii) single complexes have switch-like activation profiles where the aggregate of NEMO recruitment and time-varying properties of each complex are cytokine-specific; and, iii) that dynamics of formation and dissolution for single complexes during the primary cytokine response are independent of transcriptional feedback. Finally, we characterize a signaling motif called a variable-gain stochastic pooling network (SPN) that encompasses our experimental observations. The variable-gain SPN motif has beneficial noise-mitigation properties and provides a trade-off between information fidelity, ligand specificity, and resource allocation for intracellular signaling molecules. We propose that the variable-gain SPN architecture, and its associated benefits to signal transmission, are common mechanisms for receptor-mediated signal transduction.

## Results

### Surface receptor expression is limiting for numbers of cytokine-induced signaling complexes

IKK activity is a convergence point for pro-inflammatory signals that regulate NF-κB downstream of many cytokine receptors (*26, 30*). Ligands that bind to multiple receptors with differing kinetics (*31*), and decoy receptors that sequester or antagonize signaling complexes (*32*), layer additional complexity to signal initiation at the plasma membrane. To establish expectations for numbers and types of IKK-activating complexes, we measured surface receptor expression in U2OS cells that were previously shown to form dynamic IKK puncta in response to TNF and IL-1 (*19, 33*). Using flow cytometry with reference beads for absolute quantification, we estimated the number of surface receptors per cell for TNFR1, TNFR2, IL-1R1, IL-1R2, and IL1-R3 (Figs. 1 & S1). On average, each U2OS cell presented approximately 1300 TNFR1, 700 IL-1R1, and an abundance of IL-1R3 surface receptors. Only a small number of TNFR2 and IL-1R2 were detected on the cell surface. For reference, we measured surface receptors on HeLa and KYM1 cells (Fig. S1) and found results consistent with previous reports for TNFRs (*34-36*), and agreement with surface receptor expression in other cell lines (*37-40*)

**Figure 1:**
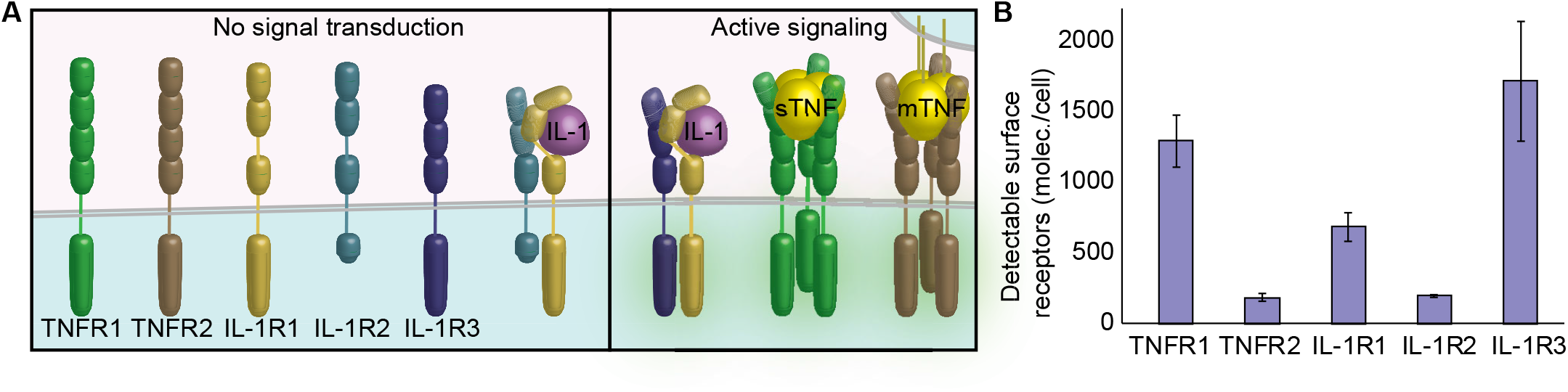
Differential expression of cytokine receptors enables cells to selectively respond to their environment. **(A**) Schematic of cognate receptors for TNF and IL-1 cytokines. Monomeric receptors and receptors that engage with decoy receptors (IL-1R2) are inactive and do not transmit signals into the cytoplasm (left). Activated receptor complexes (right), consisting of a TNFR homotrimer bound to TNF or an IL-1R1-IL-1R3 heterodimer bound to IL-1, are capable of seeding CI-like complexes in the cytoplasm. **(B)** Quantification of surface receptor expression on single U2OS cells. The average of 3-7 biological replicates is shown for each condition. Error bars represent SEM.

Although activated TNFR1 and TNFR2 both form TNF-induced homotrimeric complexes, the TNFR2 subtype binds with lower affinity to soluble TNF and shows enhanced activation by membrane-bound TNF (*31, 41*). In contrast, ligand-activated IL-1 receptor (IL-1R1) forms a heterodimer with the IL-1R3 accessory protein and dimerization can be inhibited through competitive sequestration by the IL-1R2 decoy (*32*) (Fig. 1A). Because surface expression of TNFR2 and IL-1R2 are comparably low in U2OS, receptor composition of TNF and IL-1-induced oligomers will consist predominantly of TNFR1 trimers and IL-1R1-1R3 respectively. Together with known receptor-ligand stoichiometry (Fig. 1A), our results predict that single cells can form up to hundreds of IKK-recruiting complexes for saturating cytokine concentrations (approximately 400 and 700 for TNF and IL-1 respectively). Remarkably, surface receptor expression is significantly lower than numbers for downstream signaling molecules such as NEMO that are expressed in orders of a million per cell (*42*).

### Cytokine-specific and dose-specific modulation of NEMO complex features

We set out to investigate how cytokine receptors engage NEMO as an integration hub to regulate NF-κB signaling. To counteract effects of NEMO overexpression, which can significantly inhibit NF-κB activation (Fig. S2, and (*42*)), we used CRISPR/Cas9 for targeted insertion of coding sequences for fluorescent proteins into the U2OS cell line. The resulting cells co-express N-terminal fusions EGFP-NEMO and mCh-RelA from their endogenous loci which can be used to monitor dynamic signaling events by live-cell imaging (*33*).

In response to TNF or IL-1, EGFP-NEMO transiently localizes to punctate structures near the plasma membrane (*19, 20, 33*) that are distinct from endosomal structures (Figs. 2 and S3, see also Movie S1). To further characterize the role of cytokine identity and dose on NEMO recruitment at CI-like puncta, we compared descriptive features such as their size and intensity across different cytokines and concentrations. Although properties of IL-1 and TNF-induced puncta did not show a clear trend across doses, IL-1-induced spots were significantly larger and brighter than their TNF-induced CI counterparts (Fig.2C and Movie S2; for all comparisons p-value << 10^−10^, student’s t-test). To estimate the expected number of NEMO molecules in each complex, we evaluated intensity values for each fluorescent spot in terms of a reference live-cell reporter that recruits a known number of EGFP molecules into a diffraction-limited space (*43*). By comparing cells in identical imaging conditions, our analysis suggests that each of the larger IL-1-induced spots recruit approximately 200 NEMO molecules whereas TNF-induced spots recruit around 80 (Figs.2D & S4).

**Figure 2:**
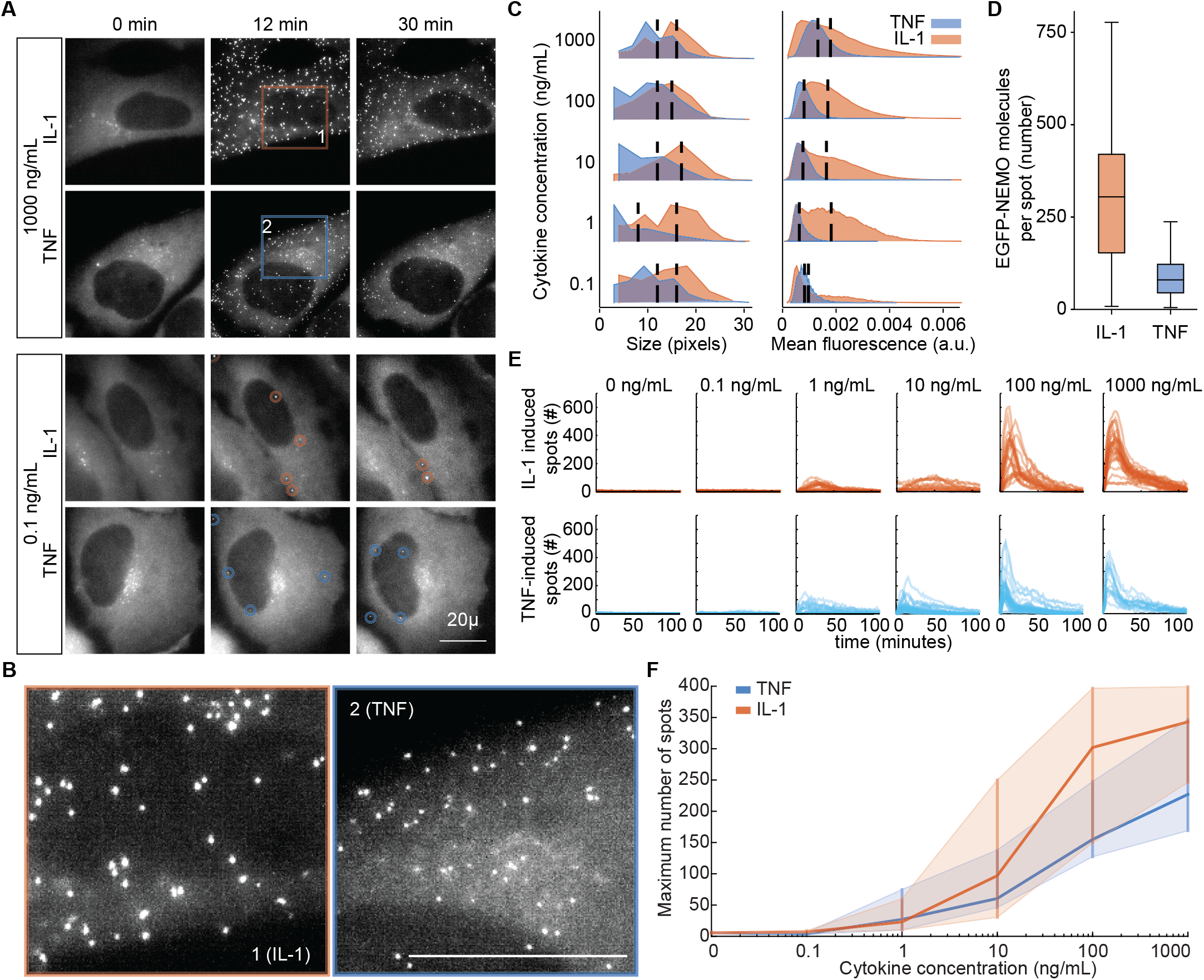
Size and intensity of NEMO complexes are modulated by cytokine identity independent of dose. **(A)** Maximum intensity projections from 3D time-lapse images of endogenous EGFP-NEMO show rapid recruitment to CI-like complexes in cells exposed to IL-1 or TNF. See also Movies S1 and S2. **(B)** Detail of fluorescent complexes (orange and blue boxes in panel A) show differences between responses to IL-1 and TNF. Scale bar represent 20µm for all. **(C)** Histograms summarizing the size (left) or intensity (right) of EGFP-NEMO complexes across concentrations of IL-1 (orange) or TNF (blue). Distributions represent single-cell data in aggregate from 3-5 experiments. Vertical bar indicates the median of each population. Comparison between all conditions, IL-1-induced complexes are larger and brighter than TNF-induced complexes (p-value << 10^−20^; t test). Numbers of cells analyzed and associated spot numbers are provided in Table S1. **(D)** Boxplot for estimates of the total number of EGFP-NEMO molecules in each CI-like complex. Median and interquartile ranges are indicated. See also methods and Fig. S2D. **(E)** Single-cell time-courses for the number of EGFP-NEMO complexes in cells exposed to indicated concentrations of IL-1 (orange) and TNF (blue). **(F)** Dose response of maximum number of EGFP-NEMO complexes. Dark orange and blue lines represents the median and vertical bars represent the interquartile range for cells stimulated with IL-1 or TNF respectively.

Time-courses for NEMO complexes in single cells showed a peak in spot numbers between 10-20 minutes and a rapid falloff thereafter (Fig. 2E), consistent with previous results from cytokine-induced IKK kinase assays (*26, 44*). Numbers of NEMO spots per cell increased with cytokine concentration and showed a tendency of higher numbers in response to IL-1 at comparable molarities (Fig.2E & 2F). Taken together, these data indicate that the size and intensity of NEMO complexes depend on the type of engaged receptor, whereas the number of complexes is modulated by cytokine dose.

### NF-κB responses are well-determined by descriptors of NEMO complexes

Next, we investigated the relationship between cytokine-induced dynamics of NEMO and RelA to ask whether NF-κB responses are well-determined by properties of fluorescent NEMO puncta when measured in the same cell. Time-courses for NEMO complexes and nuclear RelA localization were measured in response to a broad range of cytokine concentrations, and quantitative descriptors that summarize dynamic properties of EGFP-NEMO and mCh-RelA were extracted for each single cell (Fig. 3A & 3B, see also (*3, 24*)). Scatterplots of descriptors showed that cytokines and concentrations together form a continuum that relate descriptors of NEMO to RelA and correlate stronger when data are log-transformed (Figs. 3C, S5). Increased R^2^ values in log-space is likely because the range in values for descriptors of NEMO and RelA span over orders of magnitude between cells and conditions. R^2^ values for same-cell descriptors of NEMO and RelA revealed a strong effect size indicating that cell-to-cell variability in NF-κB response can be reasonably determined by descriptors of EGFP-NEMO (Fig. 3C & 3D). Multiple linear regression for combinations of NEMO descriptors only marginally increased R^2^ values (Fig. S6).

**Figure 3:**
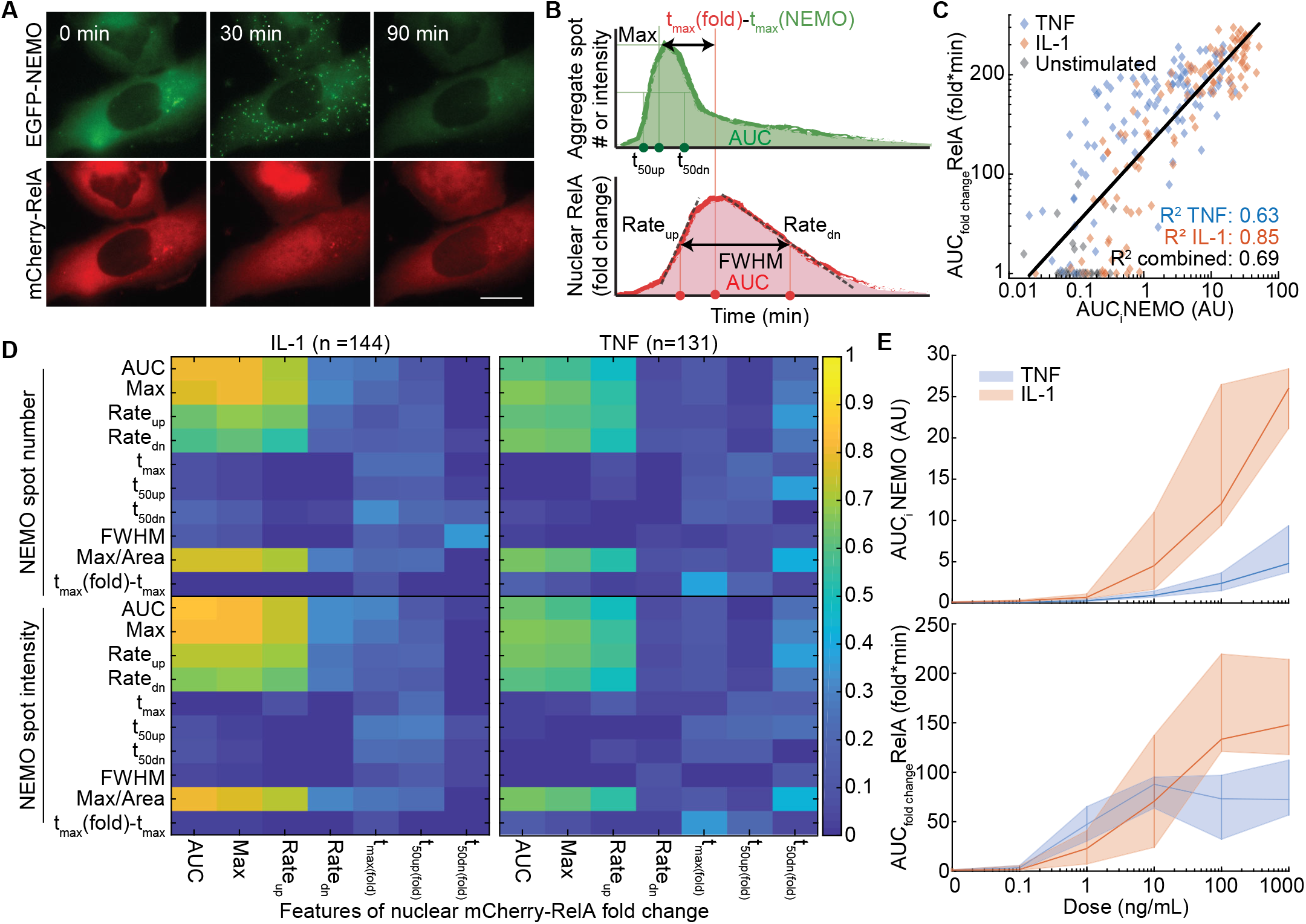
Same-cell NF-KB responses are determined by number and intensity of NEMO complexes. **(A**) Maximum intensity projections from 3D time-lapse images of EGFP-NEMO (top) and mCherry-RelA (bottom) expressed endogenously in the same U2OS cells. Cells were stimulated with 10 ng/mL IL-1. **(B)** Diagrams indicating time-course descriptors for EGFP-NEMO complexes (top) and nuclear fold-change of mCherry-RelA (bottom) in single cells. RelA features are similar to those used previously (*3, 24*). **(C)** Example of a strong same-cell correlation between single-cell descriptors of EGFP-NEMO complexes and nuclear RelA fold change. IL-1 and TNF responses across all doses overlap and form a continuum that relates the paired descriptors. **(D)** Heat map of coefficients of determination (R^2^) for all pairs of same-cell descriptors for IL-1 (left) and TNF responses (right). See also Fig. S2A and Table S2 for summary of cell numbers per condition. **(E)** Dose response curves for one of the strongest same-cell descriptor pairs. Dark orange and blue lines represents the median and vertical bars represent the interquartile range for cells stimulated with IL-1 or TNF respectively.

Previously, we showed that the ‘Area Under the Curve’ (AUC) and ‘Maximum’ (MAX) are scalar descriptors of a nuclear RelA fold-change time-course that encode the most information about cytokine dose (*3*). Notably, among all descriptors for nuclear RelA fold-change dynamics, AUC_RelA_ and MAX_RelA_ also had the strongest R^2^ with same-cell descriptors of NEMO-recruiting complexes (Fig. 3D; AUC_NEMO_ and MAX_NEMO_ respectively). Both NEMO descriptors showed similar coefficients-of-determination for RelA descriptors, whether measured in terms of numbers for EGFP-NEMO spots or aggregate intensity of EGFP-NEMO within complexes. Consistent with our previous observations for RelA descriptors, AUC_NEMO_ and MAX_NEMO_ also show dose-responsive trends that extend beyond saturating concentrations of TNF and IL-1 in terms of the NF-κB response (Figs. 2F & 3E).

Overall, same-cell measurements of EGFP-NEMO and mCh-RelA reveal that the aggregate NF-κB response in a cell is well determined by the sum of NEMO recruitment to CI-like signaling complexes near the plasma membrane. Because enrichment of NEMO at ubiquitin-rich structures is an induced-proximity mechanism for kinase activation, it’s reasonable that the AUC of EGFP-NEMO intensity in puncta is a strong proxy for downstream signaling. However, our characterization for the EGFP-NEMO fluorescence intensity at cytokine-induced spots showed wide-based distributions which could indicate that the amount of NEMO recruited at each spot varies significantly (Fig. 2C). Nevertheless, the number of EGFP-NEMO puncta (MAX_NEMO_) is almost interchangeable with AUC_NEMO_ as determinants of same-cell responses.

### Cytokine environments control numbers and timing of NEMO complex formation

Asynchronous properties of EGFP-NEMO spots, such as when a spot forms or rates for NEMO recruitment and dissolution, will contribute to variability when spot intensities are measured from a snapshot image at a single time point. To understand the extent of inter-spot and between-cell variability, we used high-frequency imaging to enable tracking of each single spot over time (Figs. 4, S7A, and Movie S3). Cells were stimulated with a step, pulse, or ramp in cytokine concentration in a microfluidic cell culture system (*45*) to observe how dynamic environments modulate properties of NEMO-recruiting complexes.

**Figure 4:**
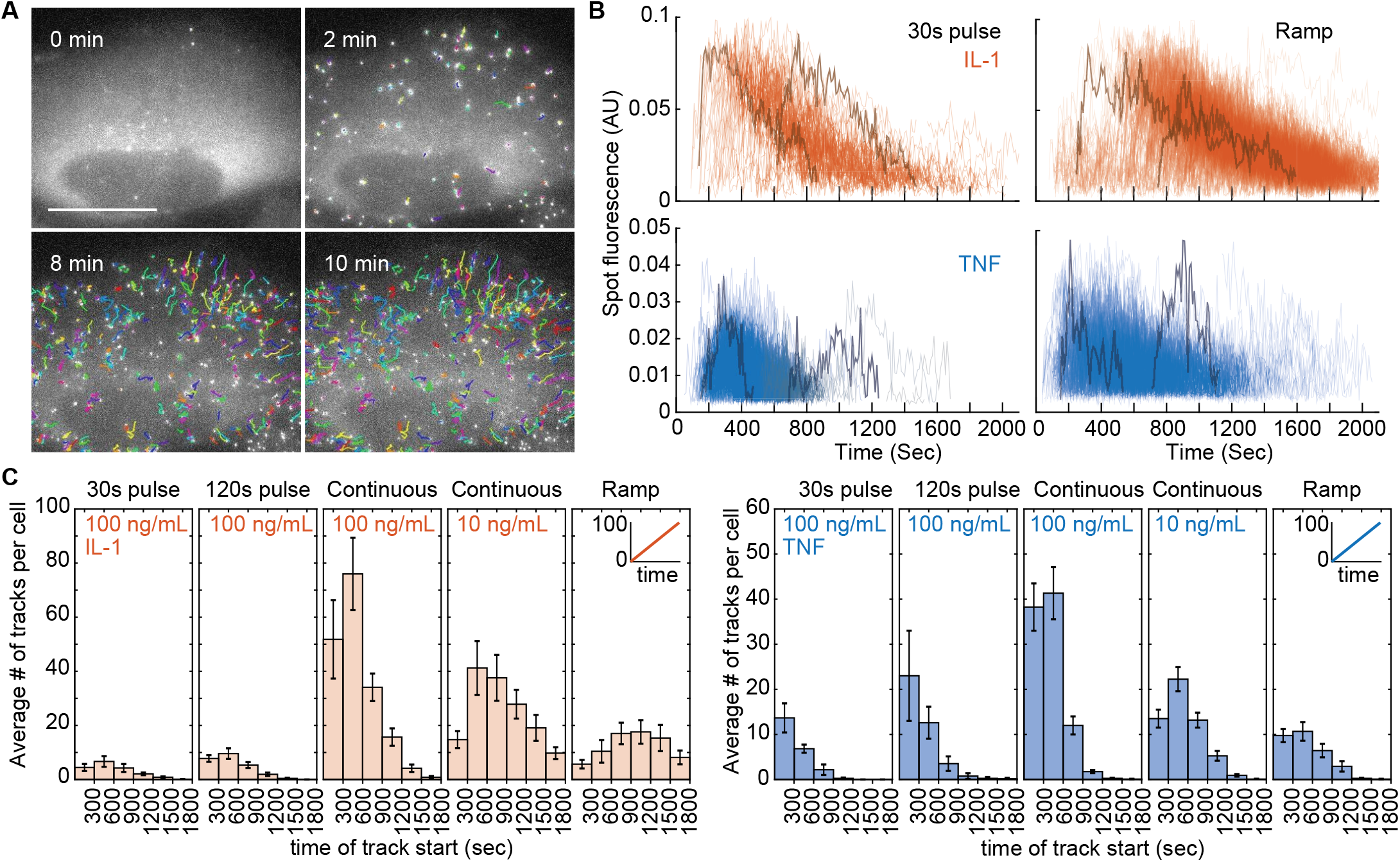
The number and time of NEMO complex formation is determined by the extracellular environment. **(A)** Maximum intensity projections from high frequency 3D time-lapse imaging experiment of cells exposed to 100 ng/mL IL-1. Colored lines in overlay indicate tracks for individual EGFP-NEMO complexes over time. On average, 60-75% of detected complexes are associated with a track of significant length (Fig. S4A). Scale bar represent 20µm, see also Movie S2. **(B)** Time-courses of fluorescence intensity for single tracked EGFP-NEMO complexes in cells stimulated with indicated conditions in a microfluidic cell culture system (*45*). Two representative single-complex trajectories are indicated in each condition (dark orange and dark blue lines). **(C)** Bar graphs for the average number and the time of formation for tracked EGFP-NEMO complexes. On average, 10 cells were analyzed in each condition. Error bars represent the SEM.

Tracking experiments revealed differences in fluorescence intensity time-courses, where IL-1 induced EGFP-NEMO puncta were consistently brighter and longer-lived than TNF-induced puncta (Fig. 4B). Both cytokines induce spots that peak within 2 to 3 minutes of detection followed by a decay phase where spots decline in intensity (Fig. 4B and S4B). When stimulated with a cytokine pulse, most spots are detected only after the cytokine is removed (Fig. 4C) demonstrating that formation of NEMO-recruiting complexes is variable and takes place up to 30 minutes following a stimulus. Step, pulse, and ramp stimulation further showed that the number of spots and the timing of spot formation are both modulated by the dynamics of cytokine presentation.

Descriptors for single spot trajectories showed remarkably low variability, in particular for the TNF response, when compared within the same cell or when the average of single-spot descriptors was compared between cells (Fig. S7C and D). Analysis of the ‘AUC spot intensity’ descriptor (AUC_i_) revealed a quadratic relationship between ‘mean AUC_i_’ and ‘AUC_i_ variance’ for spots when measured in the same cell (Fig. S7E). Increased variance between large NEMO-recruiting complexes may be due to steric properties of supramolecular assemblies. For example, where growth for certain types of ubiquitin polymers is spatially limited, or intact portions of large ubiquitin chains are clipped off en bloc or through endo-cleavage (*13*), leading to greater inter-spot variability. In nearly all cases, the coefficient of variation (CV) values for distributions of single-spot descriptors indicate that noise is lower than a Poisson processes (Fig. S7). Together, our results suggest that dynamics of NEMO-recruiting protein complexes are strictly regulated for each cytokine response.

### Negative feedback on NEMO complexes is primarily independent of transcription

NF-κB-mediated expression of A20 and subsequent deubiquitinating (DUB) activity against NEMO-recruiting complexes constitute an essential negative feedback motif in inflammatory signaling (*22*). Within tens of minutes, it’s feasible that nascent A20 contributes to the decay phase of EGFP-NEMO spot numbers observed in whole-cell measurements (Fig. 2E) thereby reducing IKK activation as observed in cell population assays (*26, 44*). It was therefore unexpected that the decay phase for single EGFP-NEMO tracks is visible within several minutes of stimulation, which is remarkably fast for a feedback mechanism governed by transcription and translation (Fig. 4B and S7).

To understand how different mechanisms of negative feedback impact trajectories of single EGFP-NEMO spots, we developed a model using ordinary differential equations. Here, cytokine stimulation induces formation of single spots at different times and each spot becomes larger and brighter with Michaelis-Menten kinetics to approximate ubiquitin polymer growth and EGFP-NEMO recruitment. The model considers two sources of negative feedback that act on NEMO-recruiting complexes by enhancing their disassembly rates. The first source aggregates the sum of NEMO-recruiting complexes to drive expression for an A20-like negative feedback mediator (‘transcriptional feedback’; Fig. 5A). The second source considers the impact from basal expression of DUBs in resting cells before stimulation (‘basal feedback’; Fig. 5A). For both sources, the strength of negative feedback on each NEMO-recruiting spot increases with size to mimic DUB recruitment to ubiquitin polymers in the complex.

**Figure 5:**
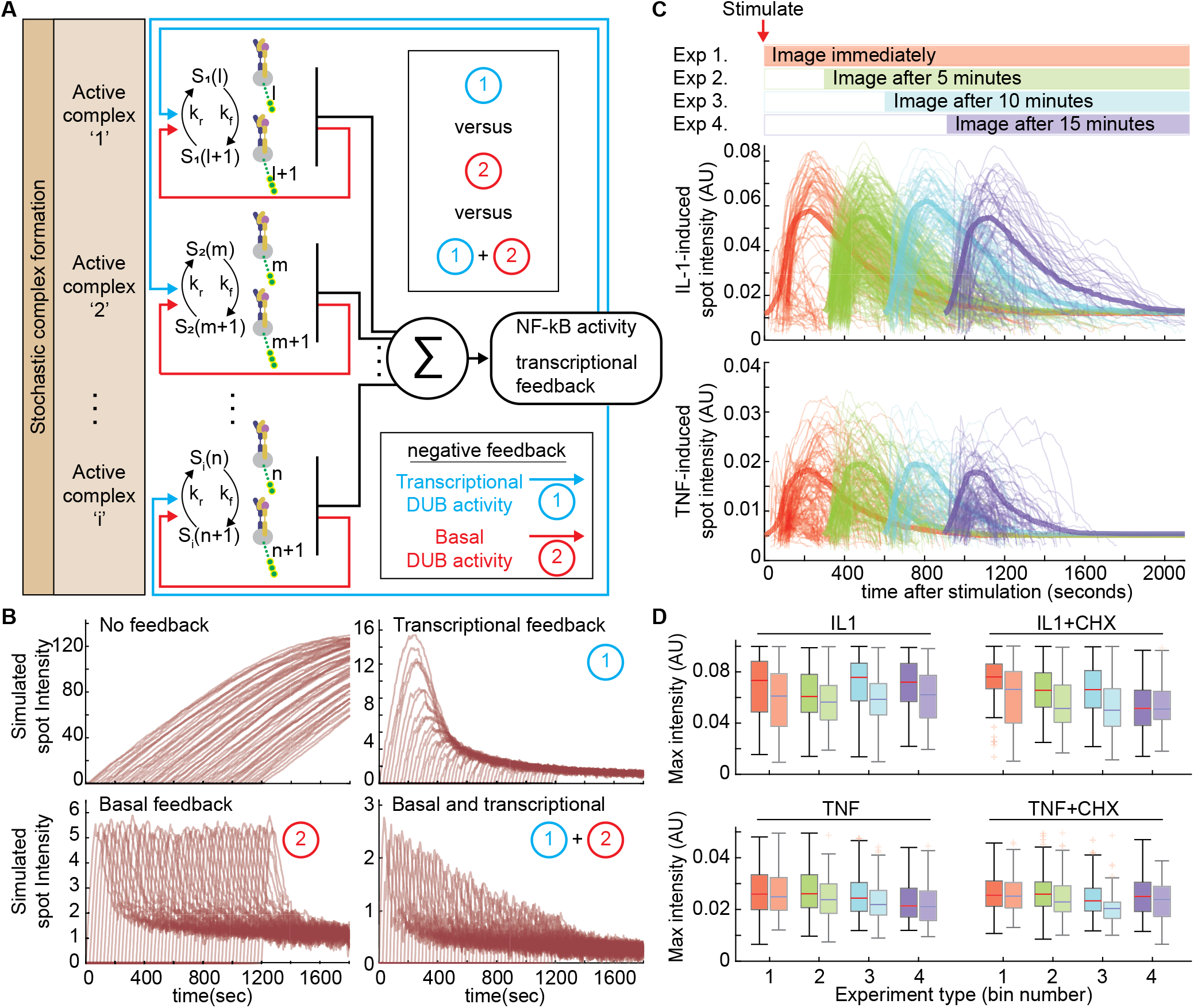
The primary NEMO response is independent of the transcriptional feedback loop. **(A)** Schematic of the hybrid deterministic−stochastic (HyDeS) model of basal (red arrows) and transcription-dependent (blue arrows) feedback regulation of NEMO complexes. Each growing CI-like complex will recede with rates that depend on recruitment of deubiquitinating enzymes (DUB). **(B)** Simulations of individual EGFP-complex trajectories using the model in fig. 5A, considering: no feedback (top left), transcription-dependent feedback only (top right), basal feedback only (bottom left), or combined transcriptional and basal feedback (bottom right). **(C)** Schematic of reverse time-course experiments. Only new EGFP-NEMO complexes that form within the first 2 minutes are tracked in movies from each high-frequency imaging experiment. Cells were stimulated with 100 ng/mL concentrations of either IL-1 (top) or TNF (bottom). **(D)** Boxplots of maximum intensity of individual EGFP-NEMO complexes trajectories after stimulation with IL-1 (top) or TNF (bottom) in cells pretreated for 20 minutes with media plus DMSO (left) or cycloheximide (CHX, right). Co-stimulation with CHX does not significantly increase values for single spot descriptors as predicted by the transcriptional feedback model (see also Fig. S5). Median and interquartile ranges are indicated, and biological replicates are shown side-by-side with increased transparency for each experiment.

Using simulations to vary the strength for each source of negative feedback, their respective impacts on single-spot dynamics was apparent (Figs. 5B and S8). By increasing the strength of basal feedback the decay phase for single-spot trajectories became steeper, and each spot displayed a sharp peak of intensity that was similar between spots regardless of when they form. By contrast, even though transcriptional feedback also increased steepness of the decay phase, the peak intensity of spots that form earlier were significantly greater than spots that formed later. For both sources, increasing strength of negative feedback reduces the overall peak height for all simulated spots (Fig. S8).

Based on simulations, if transcription is the predominant source of negative feedback, then spots that form later after cytokine stimulation are predicted to have lower maximum intensity, shorter track length, and smaller AUC (Figs. 5B and S8). To test the prediction, we performed a reverse time-course experiment where imaging started after a delay relative to the time of cytokine stimulation (Fig. 5C; 0, 5, 10, 15 minutes). Only new spots that formed within the first two minutes were tracked for each condition, thereby enabling direct comparison of early versus late-forming spots and minimizing effects of photobleaching. Biological replicates revealed that early- and late-forming single spot trajectories share highly similar dynamics (Figs. 5B, 5C, and S9), suggesting that transcriptional feedback is dispensable in regulation of dynamics for CI-like complexes. We therefore repeated reverse time-course experiments in the presence of cycloheximide (CHX) to prevent translation of NF-κB-regulated genes, effectively breaking the transcriptional negative feedback loop. CHX inhibited protein translation, evidenced by persistent nuclear NF-κB in cells co-stimulated with CHX (Fig. S10). However, loss of transcriptional feedback did not increase features of single spot trajectories as predicted by the transcriptional feedback model (Figs. 5D and S8). These results demonstrate that transcription is not the predominant mechanism of negative feedback on NEMO recruitment at CI-like complexes in the timescale of the primary cytokine response.

### Stochastic pooling mitigates noise and provides tunability between information transmission and response magnitude

For each of TNF and IL-1, trajectories of EGFP-NEMO spots are highly similar regardless of when they form during a cytokine response. This observation argues that each complex behaves as an independent switch, that when activated recruits a quantized amount of NEMO over its lifespan. Together with same-cell NEMO and RelA data, our results support a generalization of cytokine-IKK-NF-κB signaling that is evocative of a stochastic pooling network (SPN), a model sensory system with noise-mitigating and information-compressing properties (*46*). An important difference however, is that while binary detectors in an SPN transmit on-off measurements about an information source, each NEMO-recruiting complex performs amplification with a gain determined by the cytokine-receptor-complex identity.

To understand information transmission properties of the IKK-NF-κB signaling axis, we abstracted the system into parallel and independent CI-like switches that provide redundant measurements of the same extracellular signal. When a switch (CI) activates in response to a signal (S), it amplifies with a ligand-specific gain (G_L_) of NEMO activity (Fig. 6A). The total cellular response (R) in terms of NEMO activity and subsequent NF-κB translocation is the summation of all amplifier gains in a cell (Fig. 6A). We reasoned that the resulting variable-gain SPN (VG-SPN) will have different signal transduction properties, depending on the number of switches and their associated gains.

**Figure 6:**
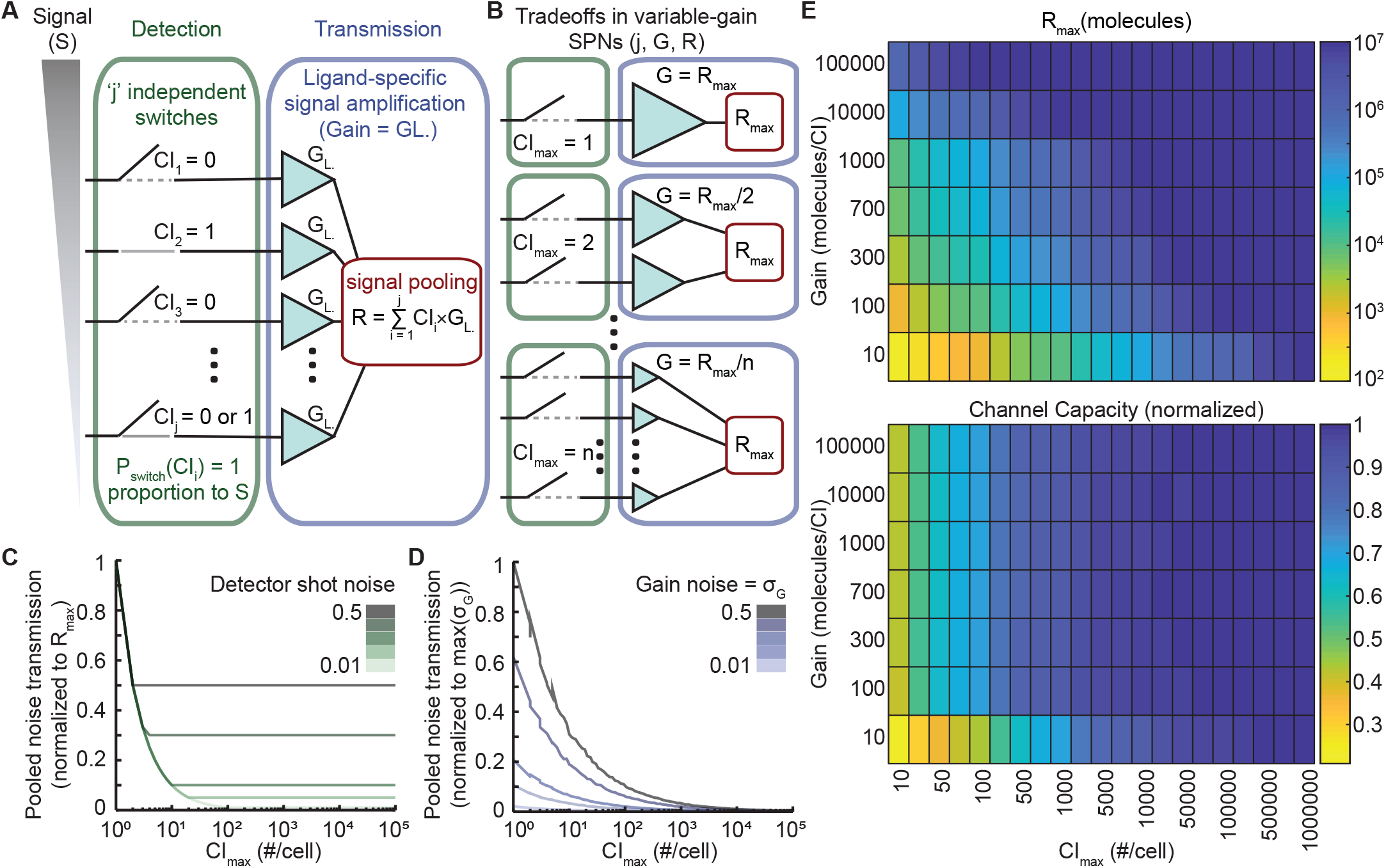
Variable-gain stochastic pooling network mitigates noise to provide efficient trade-offs with information transmission. **(A)** The VG-SPN consists of 2 steps, detection and transmission. For detection, each CI-like complex (CI) is modeled as one of ‘j’ independent switches with an activation probability proportionate to the signal strength. During transmission, each activated switch amplifies the signal with gain ‘G_L_’, related to the number of NEMO molecules recruited by each CI-like complex. NEMO from each CI-like complex is added to the cytoplasmic ‘pool’ of activated signaling molecules. **(B)** Different configurations of the VG-SPN. For a mathematically controlled comparison, CI and ‘G_L_’ are adjusted so that each configuration can achieve the same maximum response (R_max_). **(C)** Detection introduces shot noise, where a switch activates erroneously with a probability given by the shot noise level. Since each CI is independent, shot noise is mitigated through parallelization via multiple CI until a ‘noise floor’ is achieved. The noise floor is determined by the shot noise value. **(D)** Variability in the gain from each CI switch introduces noise during transmission. Gain noise is effectively averaged and approaches zero via parallelization of independent switches. **(E)** Maximum response (R_max_; upper limit was set to 10^7^ response molecules per cell) and channel capacity corresponding to different VG-SPN configurations (see also Fig. S11). The VG-SPN architecture provides ligand-specific tunability between R_max_ and information transmission.

A mathematically-controlled comparison (*28, 47*) between different network configurations was enabled by assuming each configuration is capable of producing the same maximal steady-state response (R_max_; Fig. 6B). Simulations for different mathematically-controlled configurations of the VG-SPN demonstrated that shot noise associated with signal detection, and noise associated with the signal gain, both fall off rapidly with increasing numbers of CI switches (Figs. 6C and D). Noise from these sources can be mitigated almost completely when cells are capable of forming ∼100’s-1000’s of CI per cell (CI_max_/cell). Here, noise-mitigation benefits can be attributed to signal parallelization through quantized nodes. For example, gain noise at each CI can be positive or negative, and because each CI is independent, the distribution for noise across all CI in a cell is centered near zero. Summation of signaling through R_max_ in a VG-SPN effectively averages the contribution of noise across all CI/cell, which is ideally zero assuming sufficient parallelization and absence of other biases. See also Supplementary Methods for similar description of shot noise mitigation.

We then explored the information transmission properties of VG-SPN architectures by calculating their channel capacities (*2*). We built a computational model using parameters obtained from our single-cell IKK data and simplifying assumptions about distributions for CI properties (see supplementary methods for details, and hyperparameter tuning in Fig. S11). For these simulations, CI_max_/cell and amplifier gain G_L_ were allowed to vary independently. Consistent with our previous analysis for noise-propagation in VG-SPNs, when G_L_ is greater than 10 molecules per complex the channel capacity increases with the number of CI_max_/cell and rapidly approaches saturation (Fig. 6E, bottom). However, for G_L_ values of 10 or lower, the system requires order-of-magnitude more CI_max_/cell to reach comparable channel capacity values. We also calculated the maximum response (R_max_) which shows an inverse linear relationship, where different configurations of CI_max_/cell and G_L_ can achieve activation of the same number of response molecules up to arbitrarily high numbers (Fig. 6E, top). Taken together, the VG-SPN motif generates trade-offs between surface receptor numbers, cytoplasmic molecules, and channel capacity, enabling cells to fine tune sensitivity, response magnitude, and information transmission for each ligand.

## Discussion

Endogenously expressed EGFP-NEMO is a multifaceted reporter that reveals several aspects of signal transmission. At the level of detection, there is agreement between numbers of EGFP-NEMO puncta induced by saturating cytokine concentrations and average surface receptor numbers in U2OS cells (Figs. 1 and 2). Similarly, the timing of formation of EGFP-NEMO puncta establishes when ligand-receptor-adapter assemblies become capable of signaling in the cytoplasm. Continuous stimulation experiments show that most EGFP-NEMO spots form within 5-10 minutes, indicating that cytoplasmic components of CI-like complexes are not limiting. However, in cells exposed to a short pulse, new spots form up to 30 minutes following cytokine removal, demonstrating variability in timing for receptors to assemble into a signaling-competent stoichiometry. In the cytoplasm, spot intensity time-courses inform about biochemical interactions and feedbacks linked to signal amplification. Here, families of different ubiquitin ligases, kinases, and DUBs engaged at CI-like structures establish rates of EGFP-NEMO recruitment and dissolution. Distinct properties between puncta initiated by different receptor superfamilies (Figs. 2-5) suggest that various adapters and ubiquitin requirements associated with different types of CI-like complexes will determine cytokine-specific signal amplification (*10-13, 19*). We therefore expect endogenous EGFP-NEMO will be valuable in unraveling these upstream mechanisms of inflammatory signaling, similar to reporters for NF-κB that have contributed richly for well over a decade.

Notably, imaging requirements for EGFP-NEMO limit experimental throughput, in particular for spot tracking experiments that require both high magnification and high frequency time-lapse. Although dynamical properties of NF-κB signaling are important mediators of information transmission, analyses that calculate information metrics require a large number of single-cell data points (*1-3*). Consequently, our analysis of EGFP-NEMO required simplifications through coarse-grained scalar descriptors that summarize dynamic properties of NEMO-recruiting complexes and nuclear RelA localization in the same single cells. Our analysis revealed two descriptors of EGFP-NEMO that are strong determinants for descriptors of RelA that were previously shown to carry the most information about cytokine concentrations in the milieu (*3*). As technologies emerge that enable data collection for calculation of information metrics between both reporters in the same cell, determination is likely to improve. However, it is also rational that the aggregate sum of NEMO in CI-like complexes during a primary cytokine response is a strong same-cell determinant of the accumulated NF-κB response. This deterministic relationship may therefore remain among the strongest in coming years, providing a read-out for signaling flux and perturbations at the level of CI.

Induction of A20 transcription is considered a defining negative feedback in the NF-κB response (*10, 23, 26*). However, tracking experiments revealed that trajectories of early and late-forming EGFP-NEMO spots are remarkably similar and insensitive to CHX (Fig. 5). Although this does not preclude non-catalytic roles for A20 in regulation of IKK (*29*), it demonstrates that each CI-like complex is independent and not influenced by transcriptional feedbacks from complexes that form earlier during a response in the same cell. Our data therefore support another proposed role, where A20 feedback is not primarily directed at the initial immune response but instead establishes tolerance to subsequent stimuli (*48*). Taken together, signal amplification and negative feedback at each CI-like complex is determined predominantly by the resting cell state.

Abstraction of our experimental observations revealed a VG-SPN signaling architecture (Figs. 5 and 6). The resulting model enabled us to investigate *in silico* the emergent properties of IKK-NF-κB at the level of cytokine detection and signal amplification. With detection and parallel signal amplification at independent CI nodes, our model revealed that noise is effectively mitigated with 100’s - 1000’s of signaling complexes, and greater numbers have diminishing returns per complex for information transmission. Further benefits to signal transmission through a CI amplifier allow cells to fine tune numbers of activated cytoplasmic signaling molecules, which are significantly more abundant. For NF-κB signaling, high-gain amplification enables a large repertoire of receptors to engage the same cytoplasmic pool of IKK with limited occupancy of space at the cell surface. Cells can therefore favor parsimony in receptor numbers, or increase receptor numbers with reduced amplification gain, to interface with the same size pool of signaling molecules while preserving information transmission. These trade-offs provide cells with orthogonal control points to finely tune or diversify response sensitivity to stimuli, as shown here for different inflammatory cytokines.

Our live-cell experiments revealed CI-like complexes are independent and switch-like, where each complex recruits a quantized amount of IKK over its lifespan. These were unexpected results that led to conceptual simplification and identification of the underlying signaling architecture. However, independence between detector nodes is not a required characteristic of a VG-SPN and most receptor signaling systems can therefore be viewed through a similar lens. We expect the VG-SPN is common to receptor-signaling systems, each with distinct pooling functions, feedbacks, and feedforwards that will reveal their individual trade-offs and information transmission benefits to the cell.

## Supporting information

IKK_puncta_3D

IKK_puncta_cytokines

IKK_puncta_single_trajectories

## Acknowledgements

We thank Steven Smeal, James Faeder, Anne-Ruxandra Carvunis, and Suzanne Gaudet for helpful discussions. JAC would like to dedicate this work to the memory of his mother: NGVV.

## Funding

This work was supported in part to JAC from The American Association of Immunologists Intersect Fellowship Program for Computational Scientists and Immunologists, in addition to generous funding to RECL from NIH grant R35-GM119462 and the Alfred P. Sloan Foundation.

## Author contributions

Conceptualization: JAC, CSM, RECL. Methodology: JAC, CSM, GJK, YG, SAG, DLS, QZ, RECL. Software: CSM, GJK, SAG. Formal analysis: JAC, CSM, GJK, YG, DLS. Investigation: JAC, CSM, YG, CSM, QZ. Writing – original draft: JAC, CSM, RECL. Writing – review and editing: JAC, CSM GJK, YG, SAG, DLS, QZ, RECL. Visualization: JAC, CSM, RECL. Supervision and funding acquisition: RECL.

## Data and materials availability

Data, code, and materials associated with this work will be made available upon reasonable request to RECL.

## List of Supplementary materials

Materials and Methods

Table S1 – S2

Fig S1 – S11

References (*49-55*)

Movie S1 – S3

## Notes

### Competing Interest Statement

The authors have declared no competing interest.

