## Supplementary material for "A variable-gain stochastic pooling motif mediates information transfer from receptor assemblies into NF-κB": Supp. figs & methods

**This file includes:**

Materials and Methods

Figs. S1 to S11

Movie S1 to S3 legends

Tables S1 to S2

References

### Materials and Methods

#### Reagents and Plasmids

pSpCas9n(BB)-2A-Puro (PX462) V2.0 was a gift from Feng Zhang (Addgene plasmid # 62987) (Ran, Hsu et al. 2013). The design of repair template, guide RNAs, and cloning into pSpCas9n(BB)-2A-Puro (PX462) V2.0 were described before in (Pabon, Zhang et al. 2019). The pEF-24xV4-ODC-24xPP7 plasmid for polysome quantification was a generous gift from Dr. Xiaowei Zhuang (Harvard University) (Wang, Han et al. 2016). Cycloheximide was purchased from Sigma (C4859-1ML) and used as a final concentration of 50ug/mL. Recombinant human TNF and IL-1 cytokines were purchased from Peprotech (300-01A, and 200-01B respectively).

#### Cell Culture

Parental KYM1 (female), HeLa (female), and U2OS (female) cell lines were obtained from ATCC, and HeLa cell line stably expressing scFv-GFP and tdPCP-tdTomato were a kind gift from Dr. Xiaowei Zhuang from Harvard University (Wang, Han et al. 2016). All cell lines were cultured in RPMI (KYM1), DMEM (HeLa), or McCoy's 5A (U2OS) media at 37°C and 5% $CO_2$. All media were supplemented with 10% Corning Regular FBS, 100 U/mL penicillin, 100 μg/mL streptomycin and 0.2mM L-glutamine (Invitrogen). Cells were periodically monitored for mycoplasma contamination.

#### Flow Cytometry

Fluorescence-activated cell sorting (FACS) analysis of samples and PE-beads were performed in a BD LSRFortessa machine (University of Pittsburgh-Department of

Immunology Flow cytometry facility). We used Phycoerythrin (PE)-conjugated beads (BD Biosciences, Cat. # 340495) and PE-conjugated antibodies against the following human proteins were obtained from R&D systems: TNFR1 (FAB225P), TNFR2 (FAB226P), IL1R1 (FAB269P), IL1R2 (FAB663P), IL1R3 (FAB676P), polyclonal goat IgG PE-conjugated (IC108P) and mouse IgG1 PE-conjugated (IC002P). Cells were maintained and used between 5-15 passages. For de-attaching cells from plates, we use 2 mM EDTA in cold PBS. Forty-eight hours before staining 1-2x$10^5$ cells were plated in 6-well plates. On the day of the staining cells were de-attached from the plate, transferred to polypropylene microtiter tubes (2681377, Fisherbrand™), and maintained on ice during the entire process. Fc-receptors were blocked using human BD Fc block antibody (564219) for 10-15 minutes in the dark. Next, cells were washed with 2% FBS in PBS (FACS buffer) and incubated with primary antibodies for 1 hour at 4°C in the dark. Next, samples were washed twice, and resuspended in 1 ug/mL DAPI (Cat#: D1306, Invitrogen) in FACS buffer between 0.5-1.5 hours before data acquisition. Data analysis was performed using FlowJo™ (BD, version 10.6.1_CL). Samples stained with PE isotype controls were used to set the background fluorescence by subtracting their mean fluorescence intensity (MFI) to the MFI of samples stained with specific antibodies. Next, we used the MFI from PE-conjugated beads to estimate the number of PE molecules on the surface of each cell. All these antibodies have been previously used to estimate surface receptors, with accepted 1:1 ratios of PE to antibody molecules (Serke, van Lessen et al. 1998, Bryl, Vallejo et al. 2005, Bebes, Kovács-Sólyom et al. 2014).

**Fixed-cell Immunofluorescence**

U2OS WT and U2OS cells stably overexpressing NEMO cells were seeded into plastic flat bottom 96-well imaging plates 24 hours before cytokine treatment at a density of 7000 cells/well. On the day of the experiment, wells receiving IL-1 or TNF (10 ng/mL or 100 ng/mL, respectively) were stimulated 45 min prior to fixation. Pre-warmed 15X cytokine mixture was spiked into wells and mixed. After treatment and before fixation the cells remained in environmentally controlled conditions (37°C and 5% CO2). Cells were fixed with 4% PFA and permeabilized using 100% methanol for 10 minutes each with one PBS wash in between, and three PBS-T (PBS 0.1% Tween 20) washes at the end. Next, cells were incubated in primary antibody solution (3% BSA in PBS-T) with 1 μg/mL of both α-RELA (sc-8008; Santa Cruz) and NEMO (sc-8330; Santa Cruz) at 4°C overnight. The following morning, cells were washed three times with PBS-T and incubated for 1 hour at room temperature with secondary antibody solution in 3% BSA in PBS-T (4 μg/mL of both goat anti-Mouse IgG Alexa Fluor 647 (A21235; Thermo) and goat anti-Rabbit IgG Alexa Fluor 594 (A11012; Thermo). After incubation, cells were washed twice in PBS-T and incubated in 200ng/mL Hoechst in PBS-T for 20 min. Finally, wells were washed in PBS-T and left in PBS to keep cells hydrated during imaging. Cells were imaged using Delta Vision Elite imaging system at 20x magnification with a LUCPLFLN objective (0.45NA; Olympus).

**Fixed Cell Image Analysis**

Using Cell profiler (www.cellprofiler.org, (Lamprecht, Sabatini et al. 2007)), cell nuclei were segmented using the Hoechst channel labeling DNA. Next, secondary segmentation was performed using EGFP-NEMO channel to define the cellular boundaries. The output of cell segmentation was compiled and analyzed using custom scripts in MATLAB (Mathworks, R2019b).

**Establishing EGFP-NEMO/mCherry-RELA CRISPR Double Knock-in Cells**

Single knock-in U2OS cell lines expressing EGFP-RELA/NEMO were generated previously using CRISPR-Cas9 technology as described in reference (Pabon, Zhang et al. 2019). To generate double Knock-in cells, we assembled the RELA repair template consisting of DNA sequences for a left homology arm (LHA −544 bp, chromosome 11_65663376–chromosome 11_65662383) followed by a mCherry protein coding sequence with a start codon but no stop codon and a sequence encoding 3x GGSG linker in-frame with the right homology arm (RHA +557 bp, chromosome 11_65662829–chromosome 11_65662276) from plasmids synthesized by GeneArt. Synonymous mutations were introduced to prevent interaction of the repair template and Cas9. Next, single EGFP-NEMO Knock-in U2OS cells were seeded in 6-well plates ($2 \times 10^5$ cells) and transfected next day with pSpCas9n-(BB)-2A-Puro-RELA_gRNAs and Bgl2-linearized mCherry-RelA repair template donor plasmid. A ratio of 3.5:1 FuGENE HD (Promega) to total DNA was used for transfection of 4 µg of DNA. Plasmids generation, and CRISPR modifications were all based on Ran et al. 2013 (Ran, Hsu et al. 2013).

**Live-cell Imaging**

Live cells were imaged in an environmentally controlled chamber (37 °C, 5% CO2) on a DeltaVision Elite microscope equipped with a pco.edge sCMOS camera and an Insight solid-state illumination module (GE). For detection of NEMO spots, U2OS cells expressing fluorescent protein (FP) fusions of RELA and NEMO were seeded at a density of 15,000 cells/well 24 h prior to live-cell imaging experiments on no. 1.5 glass bottom 96-well imaging plates (Matriplate). Medium was changed to phenol red-free FluoBrite Dulbecco's modified Eagle's medium (Gibco, A18967–01) between 30 min to 2 h before imaging. For detection of NEMO spots, live cells were stimulated with the indicated concentrations of TNF or IL-1. After cells adapted in the environmentally controlled chamber, we collected eight z-stack images of 0.5 µm separation in the Alexa Fluor 488 channel with an exposure of 0.04 sec and a transmission of 32%. Using these stacks of images, we reconstructed NEMO spots in 3D using image J (for display) and dNEMO (more details in "Punctate Structures Detection and Quantification" section). To test our different hypothesis, we performed two types of time-lapse imaging experiments: long-term and short-term high-frequency imaging. Images were collected over at least 3 fields per condition with a temporal resolution of 2 min per frame for long-term and 10 seconds per frame for short-term high-frequency imaging experiments. Wide-field epifluorescence and DIC images were collected using a ×60 LUCPLFLN objective. For all treatments, cytokine mixtures were prepared and prewarmed so that addition of 120 µL added to wells results in the indicated final concentration.

**Live Cell Imaging in Microfluidic Devices**

Microfluidic device fabrication and operation was done as described previously (Mokashi, Schipper et al. 2019). Briefly, dynamic stimulation devices of two inlets and one outlet were made with PDMS (Sylgard), autoclaved, washed with ethanol, and incubated with a solution of 0.002 v/v fibronectin in PBS for 24 hours at 37 °C. After incubation, excess of fibronectin was flushed with tissue culture media multiple times. Between $3x10^6$ and $8x10^6$ cells/ml were seeded by inserting 200 µl pipette tip in the outlet port. The outlet port was plugged with PDMS plugs after reaching a cell density for approximately 60% confluence. The device was incubated for at least 24 hours and PDMS plugs at the outlet were then replaced with pipette tips filled with medium. On the day of the experiment, tygon tubes (Fisher Scientific 1471139) were attached to the device with the other end connected to basins with media or cytokine treatment. The basins were then placed on corresponding platforms on the gravity pump and the attached tygon tubes were clamped. The device was then fitted to a custom adapter and placed under the microscope for imaging. The clamps on the tubes were removed at the beginning of the experiment. Cells were stimulated with desired pattern of cytokine stimulation (pulse, ramp or continuous) by changing the heights of basins by controlling corresponding stepper motors through custom codes uploaded on the Arduino mega 2560 microcontroller. The cytokine treatment was prepared with Alexa647-conjugated BSA (0.0025 v/v; Invitrogen) and imaged with CY5 filter to confirm the corresponding stimulus pattern. Images were collected under same conditions as previously described for live-cell imaging.

**Punctate Structures Detection and Quantification**

EGFP-NEMO and SunTag polysome spots were detected and quantified using our application dNEMO (Kowalczyk, Cruz et al. 2019). Briefly, dNEMO is a computational tool optimized for measurement of fluorescent puncta in fixed-cell and live-cell time-lapse images. The user-defined threshold for spot detection in dNEMO was set between 1.5

and 2.0 for all images. Reported pixel values for puncta were individually background-corrected by averaging pixels from an annular ring surrounding each spot. The width and offset for the annular ring were both set to 1 pixel. We furthermore stipulated that puncta must appear in at least 2 contiguous slices of the 3D images (out of 8 slices) to be considered valid. The same user parameters were applied to all images and single cells were manually segmented using dNEMO's keyframing function. For each single cell, spot features were measured for each new spot that formed following stimulation, yielding sets of single-cell spot features over time. NEMO puncta flat-field and background-corrected images were prepared for display using ImageJ.

**Tracking of Individual NEMO Spots**

The location and intensity data of EGFP-NEMO spots obtained with our dNEMO software (Kowalczyk, Cruz et al. 2019) was used for single-spot tracking using the uTrack package in MATLAB (Jaqaman, Loerke et al. 2008). The hyperparametrs in the uTrack package were adjusted to enhance tracking performance for TNF and IL-1 induced spots. Specifically, the time gap window was lowered to 3 frames and the gap penalty was lowered to 1 from their default values of 5 and 1.5 respectively. The minimum length of tracks segments used for gap closing was increased to 3 frames from the default of 1 frame. The merging and splitting events were considered while tracking. Properties of individual spots (intensity, size etc.) were then associated with tracking data to generate single-spot trajectories for each spot property. Trajectories lasting for less than 3 minutes were excluded from further analyses. With these settings, approximately 60% and 75% of spots respectively for TNF and IL-1 responses were tracked and included in subsequent analysis (Fig. S7A). Three main features were obtained from each single-spot trajectory:

1. Maximum intensity or $Max_i$ (peak intensity of a single-spot trajectory)
2. Integrated intensity or $AUC_i$ (sum of spot intensities at all time-points in a trajectory)
3. Track length (trajectory length i.e. time for which the spot is tracked)

Features of single-spot trajectories were then compared for dynamic stimuli experiments involving different cytokines across different cells (Fig S7); and for spots formed at different time-windows in reverse time course experiments (Fig 5D and S9B).

**Estimating Number of NEMO Molecules per NEMO Spot via Calibration with SunTag Labeled Polysomes**

To quantify the number of NEMO molecules within each NEMO complex (spot), the CRISPR labeling of NEMO with EGFP allows us to infer the number of NEMO molecules by counting GFP molecules and converting with 1:1 ratio. In counting GFP molecules in NEMO puncta, we utilized the live cell translation reporter developed by Chong Wang et. al, (Wang, Han et al. 2016) to calibrate the relation between GFP counts and measured GFP intensity and imaged it in HeLa cells with the same imaging condition used for NEMO in U2OS EGFP-NEMO cells.

During translation, one mRNA binds to multiple ribosomes simultaneously and form a large polysome complex. We assume that the positioning of each ribosome on mRNA independently satisfies a uniform distribution. The signal intensity of each fluorescence foci in the cytoplasm, representing the translating polysomes, varied depending upon the total number of ribosomes and the location of each ribosome on mRNA. When ribosomes reach the region after the coding sequence for the SunTag peptide, the intensity would be the maximum for fully assembled complex; when ribosomes are halfway towards the end, the signal would be approximately half maximal. To account for this variability, we used Monte Carlo simulations to randomly sample the positions of ribosomes and generate the distribution of numbers of SunTag peptides being produced when there are n ribosomes present on single mRNA, denoted by p_2 (m,n), where m is the number of SunTag. n, as the number of total ribosomes on each mRNA, satisfies Poisson distribution p_1 (n)=e^(-λ) λ^n/n!. The average number of ribosomes for each translation foci is measured to be 12 experimentally (Wang et al., 2016), therefore we set λ ≈ 12. The possibility of m SunTag peptides being translated on each translation foci are calculated as follows:

$$P(m) = \sum_{n=1}^{+\infty} p_2(m,n)p_1(n)$$

When n is large, p_1 (n) rapidly converges to zero. Therefore, we set n = 30 as the cutoff of the summation.

We quantified the fluorescence intensity of individual translating polysome in HeLa cells using dNEMO (Kowalczyk, Cruz et al. 2019) with the same threshold used for NEMO quantification in U2OS cells. As the GFP intensity is proportional to the number of GFP molecules, we determined the scaling factor between the measured fluorescence intensity and theoretical distribution of GFP molecule numbers. Here, we used the Nelder-Mead algorithm to minimize the square of difference between the "measured" GFP distribution and theoretical GFP distribution to obtain the scaling factor. To account for effects of photobleaching, we calculated the scaling factors independently for each frame of the time-lapse image. Using the scaling factor obtained from the translation reporter, the intensity of NEMO spots was converted to numbers of GFP molecules per spot.

#### Extracting Descriptors from NF-κB Dynamics

Descriptors of the fold-change of the FP-RELA mean intensity trajectories were extracted using custom MATLAB scripts and Image J movie explorer. The fold-change transform is carried out by dividing the FP-RELA trajectories by the initial nuclear fluorescence at the zero-time point. Those descriptors include:

- Area under the fold-change curve ($AUC_{Fold}$): a summary statistic which approximates the cell's response over time, previously found to carry the most information about a cell's response to cytokine stimulation (Zhang, Gupta et al. 2017).
- Max fold change ($F_{max}$ / $F_{initial}$): maximum value of the fold-change trajectory, previously found to determine the transcriptional response of cells stimulated with TNF (Lee, Walker et al. 2014).
- Maximal rate of nuclear entry ($Rate_{in}$): slope of line fitted to three data points between time zero and time of maximum fold change. Point along fold-change trajectory at which slope was determined was point where slope had maximal absolute value.
- Maximal rate of nuclear exit ($Rate_{out}$): slope of line fitted to three data points between time of maximum fold change and the end of the fold-change trajectory. Point along fold-change trajectory at which slope was determined was point where slope had maximal absolute value.
- Time of max ($t_{max\ (fold)}$): time at which fold-change is maximum.
- Time of half up ($t_{50up\ (fold)}$): time point at which fold-change trajectory rises to half the

max fold change.

- Time of half down ($t_{50down\ (fold)}$): time point at which fold-change trajectory falls to half the max fold change.

**Extracting Descriptors from EGFP-NEMO Dynamics**

Descriptors of trajectories for the NEMO spot number and NEMO spot intensity were extracted from single cell and single complex data using custom MATLAB scripts. The NEMO spot number trajectory is the number of NEMO spots detected per cell over time, while the NEMO spot intensity trajectory is the integrated intensity of the detected spots per cell over time. Those descriptors (per trajectory) include:

- Area under the curve (AUC): area under the trajectory integrated. For a given cell's NEMO spot number and spot intensity trajectories, this represents either the total number of spots detected, or the total intensity of the spots detected, respectively.
- Maximum value (Max): maximum number of spots / spot intensity for a given cell trajectory.
- Rate of entry ($Rate_{in}$): slope of line fitted to three data points between time zero and time of max. Point along trajectory at which slope was determined was point where slope had maximal absolute value.
- Rate of exit ($Rate_{out}$): slope of line fitted to three data points between time of max and the end of the trajectory. Point along trajectory at which slope was determined was point where slope had maximal absolute value.
- Time of max ($t_{max}$): time at which the NEMO spot trajectory trajectory is maximal.
- Time of half up ($t_{50up}$): time at which the NEMO spot trajectory rises to half the

maximum value.

- Time of half down ($t_{50down}$): time at which the NEMO spot trajectory falls to half the maximum value
- Full width at half max (FWHM): width of the peak of the NEMO spot trajectory between the time of half up and the time of half down.
- Maximum over area of the cell (Max/Area): maximum value of the NEMO spot trajectory divided by the area of the cell. The cell's area is given by the polygon output from the dNEMO results.
- Fold-change peak difference ($t_{max\ (fold)}$ - $t_{max}$): difference between the time of maximum fold change and the time of max of the NEMO spot trajectory.

**Correlation and Multiple Linear Regression of NEMO and NF-κB Descriptors**

Descriptors collected from both the FP-RELA and EGFP-NEMO trajectories were correlated in linear and log-log scales (see Supplemental Figure S5). In the absence of the pair of descriptors being correlated (e.g., an inability to compute some trajectory's $rate_{in}$ or $t_{max}$ in cells that did not respond significantly to stimulation), the single pair of descriptors were omitted from the correlation dataset, but not from the overall set of cell trajectories being analyzed. Multiple linear regression was also applied to the linear and log-transformed descriptors of the cells responding to TNF, IL-1, or the full combined experimental dataset (see Supplemental Figure S6). Linear correlation and multiple linear correlation of the descriptor sets was carried out using custom MATLAB scripts.

**Statistical Analyses**

Distributions of physical properties of NEMO puncta were compared to see if they were statistically different from one another (Figure 2C). The distributions were compared with a student's t-test (using native MATLAB functions), yielding p-values for the null hypothesis that the two distributions were from the same (normal) distribution, with identical means and standard deviations. Each null hypothesis was rejected at the 5 percent significance level with p-values $<< 10^{-20}$ for every TNF and IL-1 pair with the same dosage condition.

Linear regression was performed on NEMO predictors for the RELA responses (and the log-transformed predictors/responses) for cells responding to TNF and IL-1. Native MATLAB functions were used to calculate the $R^2$ and coefficients for the linear regression. When combining the sets of cells responding to TNF and IL-1 and correlating their features, the best $R^2$ for the combined data did not surpass a value of 0.71. Multiple linear regression was done on the NEMO predictors to see if a combination of NEMO predictor features could better explain a single RELA response feature.

Multiple linear regression was performed on NEMO predictors for RELA AUC-Fold response in cells responding to TNF and IL-1 to see whether combinations of descriptors improve $R^2$ values. $R^2$ values shown in Figure S3C are the best $R^2$ determined for the indicated number of predictors when considering all possible predictor combinations. Combination of predictors provided only marginal improvements, with maximum $R^2$ values approaching 0.7 and 0.8 for the linear and log-transformed (respectively).

**Computational Model (HyDeS) to examine relative effects of Basal and**

**Transcriptional feedback on NEMO spot dynamics**

We modeled the intensity of individual NEMO spots with a hybrid deterministic-stochastic (HyDeS) model. The purpose of this model is to examine the relative effects of basal and transcriptional feedback via DUB molecules on NEMO spot dynamics, and not necessarily to capture precisely the absolute intensity dynamics for EGFP-NEMO at each single spot. Therefore, to directly model the relative effects of basal and transcriptional feedback, we used three main variables. First, we defined the intensity of a spot (I) to represent a continuous approximation for ubiquitin chain size and NEMO activity resulting from molecules recruited at that spot. Second, we defined a variable X_basal for each individual spot as a proxy for feedback due to activity from basal DUBs expressed in resting cells. We modeled the X_basal variable such that the basal feedback increases as the corresponding NEMO spot grows through proportionate recruitment of DUB molecules on the same complex. Third, we defined a variable X_trnsxl to capture the transcriptional feedback due to the aggregate activity of all NEMO spots. Here, transcription-induced DUBs act in the same way as basal DUB molecules.

The model corresponding to N number of NEMO spots consists of 2N+1 ODEs: N for the individual spot intensities ($I_i$), N for the basal feedback variables corresponding to each spot (X_basal$_i$) and one for the transcriptional feedback variable (X_trnsxl). For each simulation, we modeled 200 single spots that form at regular intervals to approximate the response of a single cell to cytokine stimulation.

$$\frac{d[I_i]}{dt} = P_{form} * P_{bound} \frac{K_{growth}}{K_{limit} + [I_i]} - Kd_{basal} * Xlbasal_i * [I_i] - Kd_{trnsxl} * Xtrnsxl * [I_i]$$

$$\frac{d[X\_basal_i]}{dt} = Kx_{basal} * [I_i] - Kxd_{basal} * [Xbasal_i]$$

$$\frac{d[X\_trnsxl]}{dt} = Kx_{trnsxl} * \sum_i [I_i] - Kxd_{trnsxl} * [Xtrnsxl]$$

The variables and parameters in the model are as described in the following tables. We examined the relative effects of the two kinds of feedbacks by varying the parameters $Kx_{basal}$ and $Kx_{trnsxl}$, which respectively control the amount of local and global feedback, by two orders of magnitudes (Figure S8).

| Variable Name | Description |
|---|---|
| $\boldsymbol{I_i}$ | Intensity of ‘i’th spot, proxy for NEMO recruitment at that spot |
| $\boldsymbol{X\_basal_i}$ | Basal feedback variable corresponding to ‘i’th spot, proxy for feedback through basal/local DUB activity |
| $\boldsymbol{X\_trnsxl}$ | Transcriptional feedback variable, proxy for feedback through aggregate activity of all NEMO spots |

Variables used in the HyDeS model for NEMO spot intensity

| Parameter | Description | Default value |
|---|---|---|
| $\boldsymbol{P_{form}}$ | Probability that the spot is formed as a function of time. Once formed, the spot is assumed to stay active. | Modeled as uniform distribution i.e. N spots form at regular intervals during the modeling time scale. |
| $\boldsymbol{P_{bound}}$ | Probably that the ligand stays bound to the receptor and the complex is actively recruiting NEMO. | 0.9 |
| $\boldsymbol{K_{growth}}$ | Michaelis-Menton like intrinsic rate of Ub polymerization | $1.12*10^{-7}$ |
| $\boldsymbol{K_{limit}}$ | Michaelis-Menton like kinetic constant to limit intrinsic polymerization with increasing spot intensity | 0.1 |
| $\boldsymbol{Kd_{basal}}$ | Rate of basal feedback activity on | 0.032 |

| | | |
|---|---|---|
| | individual spots | |
| $Kd_{trnsxl}$ | Rate of transcriptional feedback activity on individual spots | 0.08 |
| $Kx_{basal}$ | Rate of activation of basal feedback due to Ub polymerization and DUB recruitment at the given spot | 23.3 |
| $Kxd_{basal}$ | Decay rate of basal feedback | $8.7*10^{-4}$ |
| $Kx_{trnsxl}$ | Rate of activation of transcriptional feedback due to collective activity of all NEMO spots | 0.08 |
| $Kxd_{trnsxl}$ | Decay rate of transcriptional feedback | $1.5*10^{-5}$ |

Parameters used in the HyDeS model of NEMO spot intensity

#### Variable-gain Stochastic Pooling Network model

Stochastic pooling networks (SPN) are a model sensory system in which noisy detectors are used to make independent and compressed measurements of a signal (McDonnell 2009). These measurements are then pooled to average out the uncorrelated noise and reconstruct the original signal. Here we characterize a SPN system with the decoration of variable-gain (VG-SPN). In the VG-SPN system, each detector amplifies its binary measurement with a 'Gain' before pooling. Thus, the VG-SPN can have different configurations depending on the number of detectors and the Gain per detector. In the context of cytokine signaling mediated by CI complexes, each complex acts as a binary detector of the extracellular cytokine presence and the number of NEMO molecules recruited by a complex is considered its corresponding Gain. Therefore, we define two fundamental variables in our model of VG-SPN system. First, $CI_{max}$ is the number of detectors in a VG-SPN configuration which is given by the maximum number of CI complexes that can form in a cell at saturation. Second, Gain is the number of NEMO molecules recruited at a given CI complex over the time course of its activity.

Depending on $CI_{max}$ and Gain configurations, VG-SPN systems will have different maximum response ($R_{max}$) in terms of total number of NEMO molecules recruited. To systematically examine the noise mitigation properties of different VG-SPN configurations, we used a mathematically controlled approach by assuming each configuration is capable of producing the same steady state response (Figure 6B). We therefore fixed the value of $R_{max}$ and defined the Gain per complex to be inversely proportional to the maximum number of complexes as: Gain = $R_{max}/CI_{max}$. We then varied $CI_{max}$ across orders of magnitude and calculated the noise associated with each configuration (Figure 6 C,D).

The detector shot noise is the fraction of NEMO molecules recruited when a CI complex forms erroneously i.e. in absence of the signal. We assigned the probability of a complex to erroneously form as the 'shot noise level' (SNL). Therefore, the noise propagated when the SNL is less than the uniform probability ($1/CI_{max}$) will correspond to one complex forming erroneously which, in turn, will be the Gain associated with that configuration. For example, erroneous activation of a 1-receptor system via shot noise will activate the system to its fullest capabilities. Due to the inverse correlation of $CI_{max}$ with Gain, the shot noise will decrease with $CI_{max}$ (Figure 6C). However, after reaching a certain value of $CI_{max}$, the SNL will become higher than the uniform probability allowing more than one complexes to erroneously form at the same time. Configurations beyond such value of $CI_{max}$ will hit the noise floor and the shot noise will not decrease further.

$$Fractional\ Shot\ Noise = \left\{ \begin{array}{ll} \frac{Gain}{R_{max}} = \frac{1}{CI_{max}}, & when\ SNL < \frac{1}{CI_{max}} \\ \frac{SNL * CI_{max} * Gain}{R_{max}} = SNL, & when\ SNL > \frac{1}{CI_{max}} \end{array} \right\}$$

We observed considerable variability in intensities of NEMO complexes formed in response to a particular cytokine (Figures 2F, 5C,D). This variability suggests that although the mean value of Gain per complex is cytokine specific, the gain associated with a single complex can vary among different complexes within a cell, and these differences will introduce ‘Gain noise’ in the pooled response. Specifically, Gain noise captures the noise in the pooled response corresponding to same number of active detector complexes due the variability in Gain per complex (i.e. the number of NEMO molecules recruited to a single complex over its lifespan). We examined the effect of different VG-SPN configurations on the Gain noise using the mathematically controlled approached as described earlier. We modeled the Gain per complex with Gaussian distribution and using the coefficient of variation given by the ‘Gain noise level’ (GNL) as: $\sigma_G$ = GNL*Gain. Therefore, for a given VG-SPN configuration, each cell will have a response corresponding to the sum of $CI_{max}$ random numbers obtained from the distribution $\mathcal{N}(Gain, \sigma_G)$. We simulated 1000 cells for every configuration with different GNL values and calculated the Gain noise (coefficient of variation) that contribute to the resulting responses (Figure 6D).

Next, we developed a computational VG-SPN model to generate simulated data for channel capacity calculations using the framework developed by (Selimkhanov, Taylor et al. 2014) and codes from our previous study (Zhang, Gupta et al. 2017). In this model, the signal S in the range (0,1] represents the extracellular cytokine dose where S=1 corresponds to saturating dose. The dose range is equally divided in discrete levels controlled by the hyperparameter ‘numDoses’.

$$S_i \in (0,1]\,, i \in [1, numDoses]$$

For each input signal $S_i$, we simulate responses from N cells, where N is another hyperparameter. The response of jth cell to ith level of input signal ($R_{ij}$) is a scalar quantity representing the total number of recruited NEMO molecules. The model thus produces simulated data as response vectors of length N for each of the 'numDoses' number of input conditions. The data workflow is as follows:

$$S = \begin{bmatrix} S_1 \\ S_2 \\ \vdots \\ S_i \\ \vdots \\ S_{numDoses} \end{bmatrix} \rightarrow R = \begin{bmatrix} R_1 \\ R_2 \\ \vdots \\ R_i \\ \vdots \\ R_{numDoses} \end{bmatrix} where\ S_i \rightarrow R_i = [R_{i1} \quad R_{i2} \quad \dots \quad R_{ij} \quad \dots \quad R_{iN}]$$

As described earlier, $CI_{max}$ is the maximum number of NEMO complexes a cell can form at saturating conditions i.e. when $S_i$ = 1. $CI_{max}$ will be related to the number of surface receptors (SR) per cell through the cytokine-dependent valency as: $CI_{max}$ = SR/valency. Therefore, to model cell-to-cell variability in receptor expression (Figure 1B), we assumed that $CI_{max}$ follows a Gaussian distribution over the N simulated cells with coefficient of variation given by the Receptor Noise Level (RNL).

$$CI_{max_j} \sim \mathcal{N}(CI_{max}, RNL * CI_{max})$$

Here the mean $CI_{max}$ is set by the VG-SPN configuration and RNL is another hyperparameter controlling the coefficient of variation in receptor expression. For instance, for N=1000, $CI_{max}$=100 and RNL=0.3, we sample 1000 normally distributed integer random numbers from $\mathcal{N}(100,30)$ to simulate responses for 1000 cells where each cell can maximally form around 100 complexes with coefficient of variation 0.3.

We used binomial distribution to model switch-like properties of complex formation in the detection step (Figure 6A) wherein the probability of complex formation ($P_{form}$) increases with the input signal $S_i$. We used modified Hill function to relate $S_i$ with $P_{form}$ (Figure 2E).

$$CI^*_{max_j} \sim Binomial\left(CI_{max_j}, P_{form}\right), \qquad where \quad P_{form} = \frac{(1+0.5) * S_i^3}{0.5 + S_i^3}$$

Here $CI^*_{max_j}$ is the number of complexes formed in response to the dose level $S_i$ in a cell having capacity to form $CI_{max_j}$ complexes at saturation. Therefore, for a given 'j'th cell, we generate $CI_{max_j}$ random numbers between [0,1]. We then check how many of those fall below the $P_{form}$ value corresponding to the given signal $S_i$. This number, $CI^*_{max_j}$, will be used as number of active complexes in the 'j'th cell. Thus, the output of detection step corresponding to signal $S_i$ is the following vector:

$$S_i \rightarrow \left[CI^*_{max_1} \quad CI^*_{max_2} \quad \ldots \quad CI^*_{max_j} \quad \ldots \quad CI^*_{max_{N^\#}}\right]$$

Note that $N^\# \leq N$ because for lower doses there might be some cells that don't cross the $P_{form}$ threshold i.e. they don't form any spots.

In the detection step, we simulated the number of complexes formed by a given cell ($CI^*_{max_j}$) in response to a given dose ($S_i$). Next, in the transmission step (Figure 6A), we model the Gain ($G_k$) in terms of number of NEMO molecules recruited per complex. The response of a cell i.e. total number of recruited NEMO molecules is then sum of the Gains associated with all the complexes formed in that cell.

$$Response\ R\left(S_i, CI_{max_j}\right) = \sum_{k=1}^{CI^*_{max_j}} G_k$$

We assume a linear relation between Gain per complex (in terms of NEMO molecules) and the experimentally observed integrated intensity ($AUC_i$, Figure S5).

$$G_k(NEMO\ Molecules) = AUCi\_to\_NEMO\_factor * \ G_k(Intensity)$$

Here AUCi_to_NEMO_factor is another hyperparameter. As previously stated for calculation of Gain noise, we use Gaussian distribution to model the variability in $G_k$ within a cell.

$$G_k \sim \mathcal{N}(Gain, GNL * Gain)$$

Here GNL is the Gain Noise Level i.e. the coefficient of variation of Gains corresponding to different complexes within a cell. We use the quadratic relation between intracellular mean and variance of $AUC_i$ (Figure S4E) to get GNL for different mean values of Gain corresponding to different VG-SPN configurations.

To summarize, response of ‘j’th cell with a VG-SPN configuration given by ($CI_{max}$, Gain) to the input signal level $S_i$ is calculated as following:

$$Response\ R_{ij}(S_i, CI_{max}, Gain) = \sum_{k=1}^{CI^*_{max_j}} AUCi\_to\_NEMO\_factor * \mathcal{N}(Gain, GNL * Gain)_k$$

$$Where \qquad CI^*_{max_j} = Count(CI_{max_j} < P_{form}(S_i))$$

$$And \qquad CI_{max_j} \sim \mathcal{N}(CI_{max}, RNL * CI_{max})$$

This process is repeated for N number of cells and ‘numDoses’ number of input signal levels:

$$CI_{max_j} \in \begin{bmatrix} CI_{max_1} & CI_{max_2} & \cdots & CI_{max_N} \end{bmatrix}$$

$$S_i \in [S_1 \quad S_2 \quad \cdots \quad S_{numDoses}]$$

There are two independent variables in this model, $CI_{max}$ and Gain, which completely define the VG-SPN configuration. In other words, the model allows independent ‘tuning’ of maximum number of NEMO complexes (i.e. number of surface receptors) and the Gain (i.e. number of NEMO recruited by each complex) to achieve a certain channel capacity.

The model has four hyperparameters: numDoses (controlling number of discrete dose levels), N (number of cells to simulate per dose level), RNL (coefficient of variation in $CI_{max}$ over cells) and AUCi_to_NEMO_factor (relating integrated intensity to number of NEMO molecules per complex). In addition to these four, the channel capacity calculation framework developed by (Selimkhanov, Taylor et al. 2014) requires another hyperparameter 'K' which controls the K-nn classification to estimate probability densities of response given input dose level. To interface the VG-SPN model with our previous code for channel capacity calculations (Zhang, Gupta et al. 2017), we adjusted these five hyperparameters to avoid technical biases that can artificially limit calculated channel capacity values for a VG-SPN configuration.

After running preliminary computational experiments, we arrived at a set of values for the five hyperparameters. Then, for each hyperparameter, we varied its value while keeping other values in the set constant to confirm that the chosen value doesn't limit the maximum channel capacity value of a VG-SPN configuration. By iterating through this process several times, and repeating it for multiple values of Rmax, we converged on the following settings (Fig S11A).

First, the number of simulated input conditions can artificially limit the channel capacity calculated for a signaling system, but becomes increasingly costly to compute when input conditions become excessively high. We simulated our model with 10, 30, 60 and 100 input conditions ('numDoses' parameter in the methods) and found that the max channel capacity does not increase beyond 60. Next, we observed that the max channel capacity decreases as the cell-to-cell variability in number of CI per cell (RNL parameter) increases. We therefore chose the intermediate value of stdR = 0.3 that is consistent with our FACS receptor analysis (Figs. 1 and S1). Next, the value of K used for K-NN classification introduces minial bias beyond a value of 5 and contributes to computational cost at higher values. We therefore chose K = 5, consistent with previous calculations (Selimkhanov, Taylor et al. 2014). We then found that simulating more than 1000 cells per input condition does not increase the channel capacity but significantly affects simulation time, so we chose N = 1000. Finally, in order to order to relate amplifier gain ($G_L$) to the quadratic noise relationship observed in data (Fig. S7E), we estimated the factor to convert integrated spot intensity to number of NEMO molecules at a CI-like complex ('AUCi_to_NEMO_factor'). Values for the conversion factor did not have significant effects on max channel capacity beyond a value of 200.

Once we fixed the hyperparameters of the model, we removed the constraints of the mathematically controlled framework and allowed the two model variables, $CI_{max}$ and Gain, to vary independently. We then calculated the channel capacities corresponding to VG-SPN configurations given by different ($CI_{max}$, Gain) pairs using previous codes (Zhang, Gupta et al. 2017) which are based on the framework developed by (Selimkhanov, Taylor et al. 2014). We also calculated the maximum response ($R_{max}$) corresponding to each ($CI_{max}$, G) configuration which is simply given by $CI_{max}$*Gain. Effectively, the value of $R_{max}$ relates to the number of NEMO or other response molecules in the cytoplasm that can be activated by a VG-SPN configuration.

| **Variable** | **Description** | **Formula** |
|---|---|---|
| $CI_{max}$ | Maximum number of CI complexes that a cell can form at saturation | Independent variable |
| Gain | Number of NEMO molecules recruited by a CI complex | Independent variable |
| $\boldsymbol{CI_{max_j}}$ | $CI_{max}$ corresponding to 'j'th cell | $CI_{max_j} \sim \mathcal{N}(CI_{max}, RNL * CI_{max})$ |

| | | |
|---|---|---|
| $P_{form}$ | Probability of complex formation as a function of input dose level $S_i$ | $P_{form} = \frac{(1+0.5) * S_i^3}{0.5 + S_i^3}$ |
| $\boldsymbol{CI^*_{max_j}}$ | Number of CI complexes formed in response to a given input dose level in 'j'th cell | $CI^*_{max_j} \sim Binomial\left(CI_{max_j}, P_{form}\right)$ |
| $G_k$ | Gain associated with 'k'th complex within a cell | $G_k \sim \mathcal{N}(Gain, GNL * Gain)$ |
| GNL | Coefficient of variation of Gains associated with different CI complexes within a cell | GNL = 0.495*Gain$^2$ – 0.096*Gain + 0.05 (where Gain is in terms of $AUC_i$) |

Variables used in the VG-SPN model

| Hyperparameter | Description | Value |
|---|---|---|
| numDoses | Number of discrete input dose levels used to simulate the model response | 60 |
| N | Number of cells simulated for each input dose level | 1000 |
| RNL | Receptor noise level in terms of coefficient of variation in $CI_{max}$ across cells | 0.3 |
| $AUC_i$_to_NEMO_factor | Factor to convert integrated intensity (AUCi) of observed CI spots into number of NEMO molecules recruited at that complex | 200 |
| K | Value of K used for K-nn classification in the framework developed by (Selimkhanov, Taylor et al. 2014) | 5 |

Hyperparameters used in the VG-SPN model

#### Supplementary figures

**Figure S1: Quantification of surface receptors in U2OS, HeLa, and KYM-1 cells.**

**(A)** Flow-activated cell sorting (FACS) distributions for surface receptor expression. Unstimulated U2OS cells were stained for with PE-conjugated antibodies against indicated surface receptors. Semi-transparent histogram represent cells stained with negative isotype control antibodies. Surface receptor numbers were subsequently calculated using BD Quantibrite™ PE beads as standard curve. **(B)** Quantification of surface expression for TNFR1, TNFR2, and IL-1R1 was performed similarly with HeLa and KYM-1 cells for comparison with U2OS data and previously published results. The average of 3-7 biological replicates is shown for each condition. Error bars represent SEM.

Fig. S1

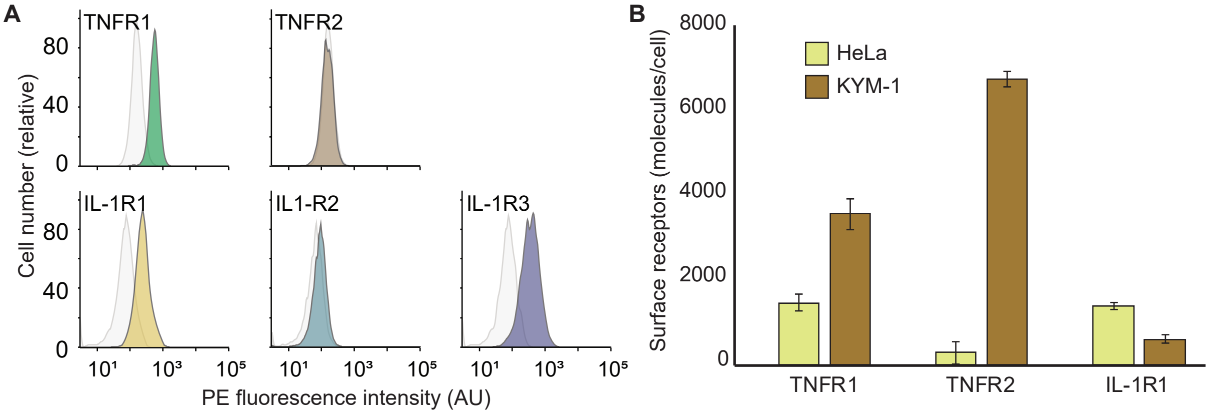

A
TNFR1
TNFR2
IL-1R1
IL1-R2
IL-1R3
Cell number (relative)
80
40
0
10¹
10³
10⁵
PE fluorescence intensity (AU)
B
Surface receptors (molecules/cell)
8000
6000
4000
2000
0
HeLa
KYM-1
TNFR1
TNFR2
IL-1R1

**Figure S2: Overexpression of NEMO interferes with the downstream NF-kB response.**

**(A)** Fixed-cell images of U2OS cells stained with fluorescent antibodies against RelA and NEMO. Hoechst counterstain was used to identify nuclei. U2OS wild-type cells (WT) or U2OS stably transfected to overexpress NEMO were stimulated with TNF or IL-1 and translocation of RelA (NF-kB) was quantified in nuclei. For stable NEMO overexpression, two cell subpopulations were visible, one with near endogenous levels of expression and the other with high expression (marked with an orange perimeter). Scale bar represent 20µm for all. **(B)** Nuclear RelA abundance from fixed-cell images described in (A) using measurements from approximately 1000 single cells per condition. For both WT and NEMO-low cells, clear accumulation of nuclear RelA is observed in response to TNF and IL-1 ($p < 10^{15}$; 1-tailed t test). However, in NEMO-high cells basal nuclear RelA was increased and cytokine stimulation did not induce nuclear translocation of RelA.

Fig. S2

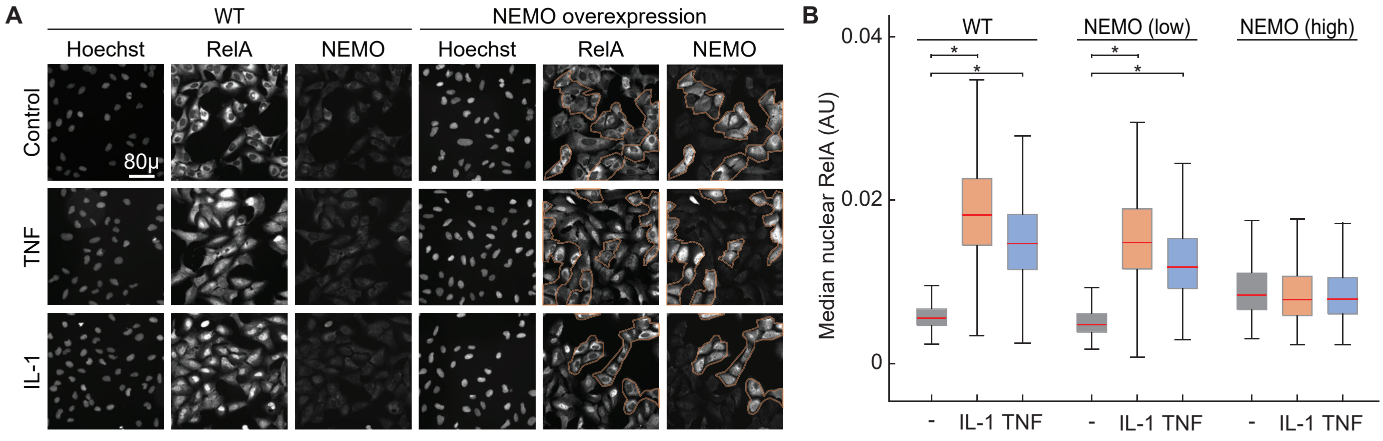

A
WT
NEMO overexpression
Hoechst
RelA
NEMO
Hoechst
RelA
NEMO
Control
TNF
IL-1
80μ
B
WT
NEMO (low)
NEMO (high)
Median nuclear RelA (AU)
0.04
0.02
0
-
IL-1
TNF

**Figure S3: Endogenous fusion EGFP-NEMO proteins do not colocalize with lysosomal structures.**

U2OS cells with endogenous EGFP-NEMO were preloaded with LysoTracker™ Blue DND-22 (ThermoFisher) and stimulated with IL-1 or TNF. Maximum intensity projections of 3D images demonstrate that EGFP-NEMO complexes do not co-localize with labeled lysosomes. Scale bar represents 20µm.

Fig. S3

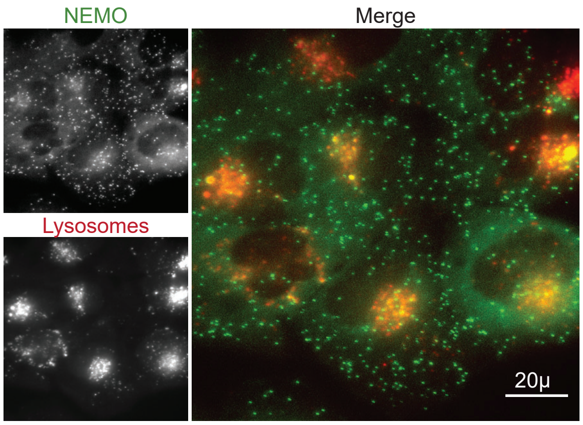

**Figure S4: Polysome calibration to estimate numbers of EGFP-NEMO molecules per complex.**

U2OS EGFP-NEMO cells stimulated with 100ng/mL IL-1 or TNF were imaged by time-lapse microscopy. In parallel, HeLa cells expressing scFv-EGFP suntag and transfected with V4-PP7 to reveal polysomes (cells were provided by the Zhuang group and prepared as described in their original work (Wang, Han et al. 2016)). Imaging conditions were identical for all. Polysome images, with a known number of EGFP molecules per spot were then used to calibrate the intensity of EGFP-NEMO complexes in TNF- or IL-1 stimulated cells at a time when cytokine-induced complexes are brightest (red line). The comparison enabled an estimation for the number of EGFP-NEMO molecules per CI-like complex. Representative maximum intensity projection images are shown for each. Scale bar represents 20µm.

Fig. S4

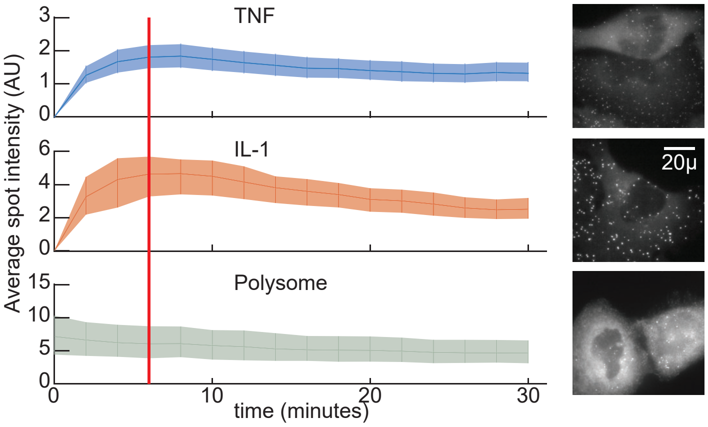

TNF
IL-1
Polysome
Average spot intensity (AU)
time (minutes)
0
10
20
30
3
2
1
0
6
4
2
0
15
10
5
0
20μ

**Figure S5: Same-cell coefficients of determination for descriptors of EGFP-NEMO and mCh-RelA dynamics.**

**(A)** Linear regression of descriptors of NEMO spot numbers (left) and aggregate spot intensities (right) against the area under the curve of the RelA fold change curve ($AUC_{fold}$). NEMO descriptors in addition to those described in Fig. 3 in the main article include the Full Width Half Maximum (FWHM) in addition to descriptors normalized to the cell area. Although correlations are shown only against the $AUC_{fold}$, each NEMO descriptor was calculated against each descriptor of RelA as shown in Fig. 3 of the main article. (B) For comparison, coefficients of determination were calculated for log-transformed data. The descriptors with the strongest coefficients of determination from linear data showed consistent improvement when data are log-transformed.

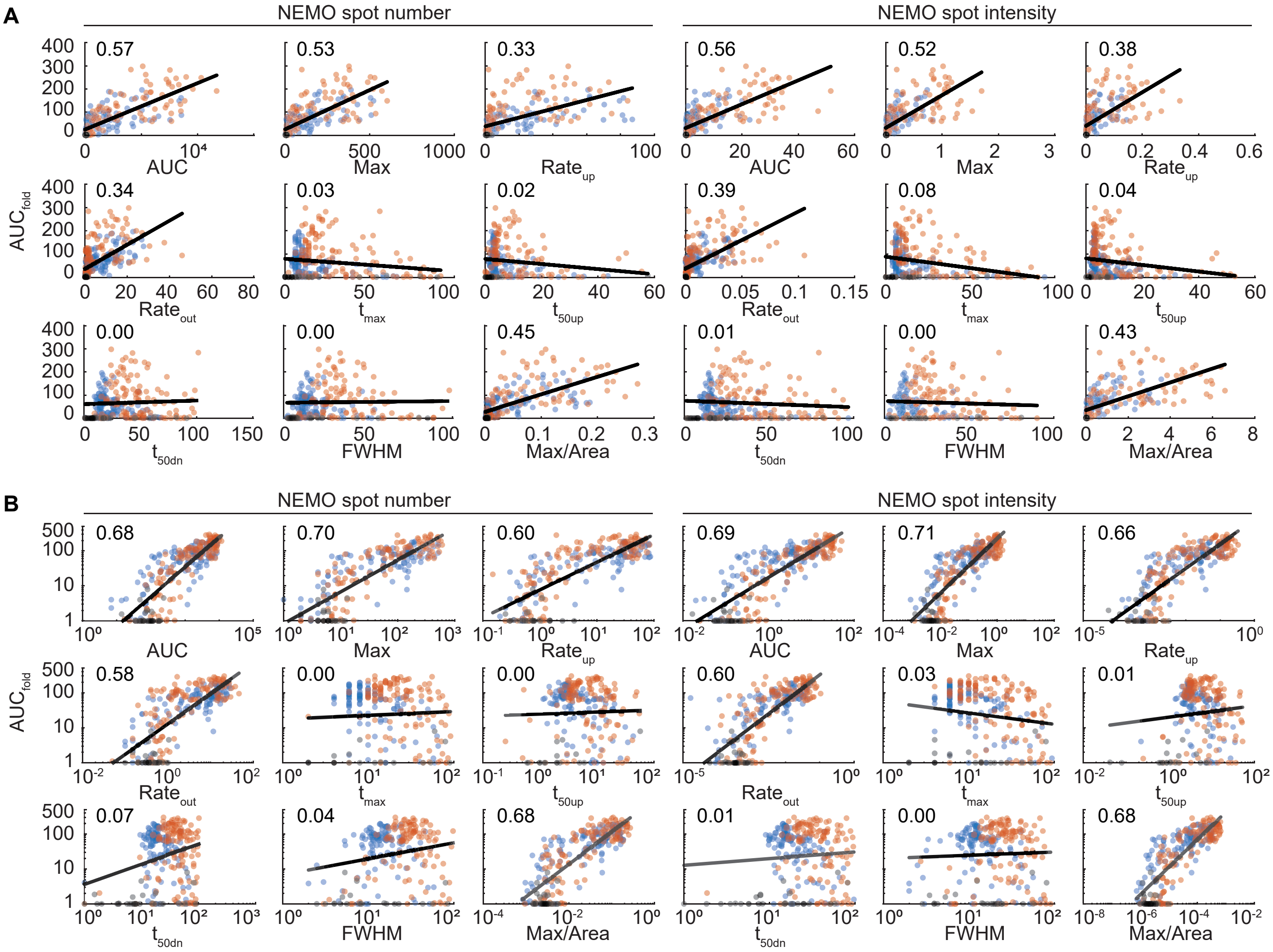
A
NEMO spot number
NEMO spot intensity
AUC_fold
0.57
0.53
0.33
0.56
0.52
0.38
AUC
Max
Rate_up
0.34
0.03
0.02
0.39
0.08
0.04
Rate_out
t_max
t_50up
0.00
0.00
0.45
0.01
0.00
0.43
t_50dn
FWHM
Max/Area
B
NEMO spot number
NEMO spot intensity
0.68
0.70
0.60
0.69
0.71
0.66
0.58
0.00
0.00
0.60
0.03
0.01
0.07
0.04
0.68
0.01
0.00
0.68

**Figure S6: Multiple linear regression do not greatly increase determination between EGFP-NEMO and mCh-RelA.**

The maximum $R^2$ value for multiple linear regression analysis for descriptors of EGFP-NEMO complexes against the $AUC_{fold}$ of nuclear RelA. For untransformed linear data (left), multiple linear regression increases $R^2$ values up to 7 predictors. However, even with 7 predictors, the $R^2$ value is still lower than all $R^2$ values for log-transformed data (right). For log-transformed data, inclusion of a second predictor showed modest improvements in $R^2$ values. Beyond the second predictor, further improvements were marginal or non-existent.

Fig. S6

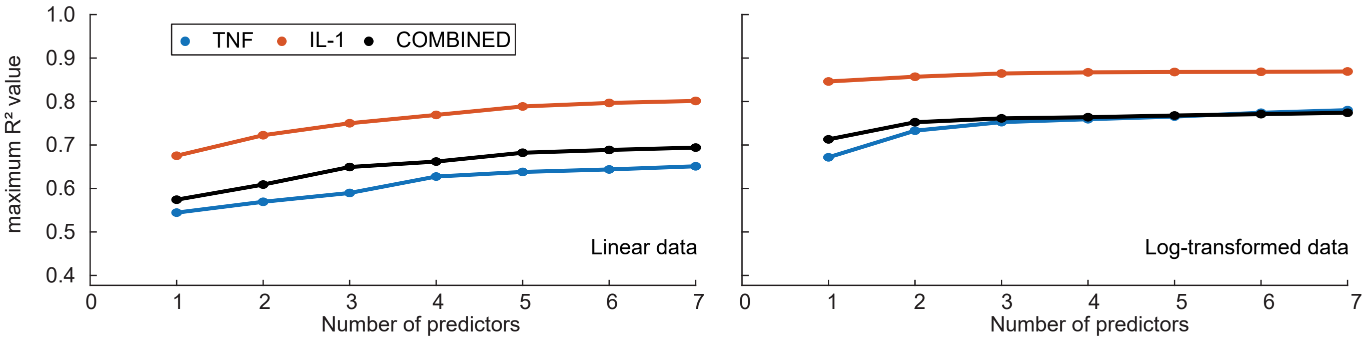

TNF
IL-1
COMBINED
maximum R² value
1.0
0.9
0.8
0.7
0.6
0.5
0.4
Linear data
Log-transformed data
0 1 2 3 4 5 6 7
Number of predictors
0 1 2 3 4 5 6 7
Number of predictors

**Figure S7**: **Tracked spots and their intra- and inter-cellular variability.**

**(A)** Boxplots (median and inter-quartile range) for the fraction of detected spots that belong to tracks across single cells. All experimental conditions were combined for each of TNF and IL-1. This analysis shows that approximately 60% of TNF-induced complexes and 75% of IL1-induced complexes were represented by single-complex trajectories from tracking experiments. **(B)** Schematic of descriptors for single-complex trajectories include maximum intensity ($Max_i$), area under the curve of intensity ($AUC_i$), and time of peak ($\Delta t$). The time of peak ($\Delta t$) is within three minutes of complex formation for both TNF and IL1 responses. **(C)** Coefficients of variation were calculated for descriptors of all complex trajectories detected within the same cell. Boxplots are distributions of single-cell CV values among all cells for each of TNF and IL1 responses. These data indicate that there is more within-cell variability for descriptors for IL1-induced complexes when compared with complexes formed in response to TNF. **(D)** The mean of descriptor values for all complex trajectories in a cell were calculated. The tight distributions for each of the TNF and IL-1-induced complexes (CV values are inset) indicate that the average of descriptors for single complex trajectories are remarkably similar between single cells. For both $AUC_i$ and $Max_i$, descriptors are 3-4 times lower for TNF-induced spots than IL1-induced spots, which is consistent with biophysical differences between cytokine-induced spots described in Fig. 2 of the main article. **(E)** For single cells, the plot between mean and variance of $AUC_i$ of all spots within a cell follows a quadratic relation. This result suggests that as single complexes become larger and longer-lived in a cell, the corresponding variance between single spots in a cell increases multiplicatively. In general, both TNF and IL1 treated cells fall below the y=x line indicating sub-Poisson noise.

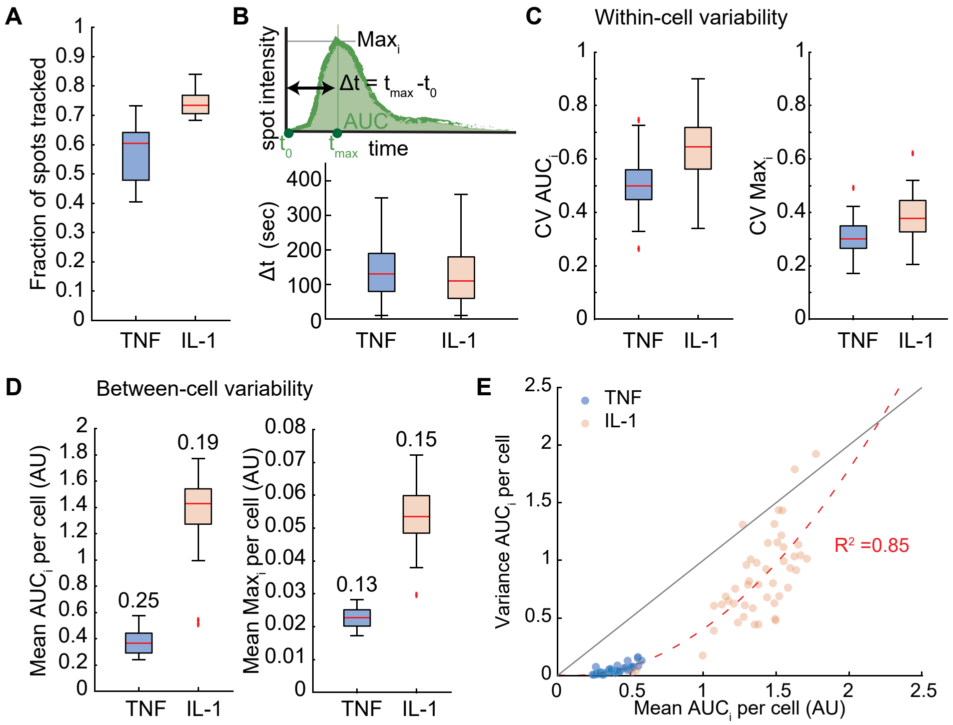
A
B
C
Within-cell variability
D
Between-cell variability
E
Fraction of spots tracked
TNF
IL-1
spot intensity
Max_i
Δt = t_max -t_0
AUC
t_0
t_max
time
Δt (sec)
CV AUC_i
CV Max_i
Mean AUC_i per cell (AU)
0.25
0.19
Mean Max_i per cell (AU)
0.13
0.15
Variance AUC_i per cell
Mean AUC_i per cell (AU)
R² =0.85

**Figure S8: Simulations of transcriptional and basal negative feedback on single-complex NEMO time courses.**

Matrix of simulations of individual EGFP-complex trajectories using the HyDeS model in Fig. 5A. Parameter sweeps for increasing strength of transcriptional feedback (left to right) and increasing basal feedback (top to bottom) were used to observe trends in single-complex trajectories that form in regular intervals after cytokine stimulation. For both sources of negative feedback, increasing feedback strength reduces the maximum intensity and AUC of each complex. Negative feedback through transcription uniquely causes late-forming complex trajectories with maximum intensity and AUC that are lower than early-forming complexes in the same simulation. Although basal negative feedback alters properties such as the sharpness of a trajectory, it operates identically on early- and late-forming complexes in the same simulation.

Fig. S8

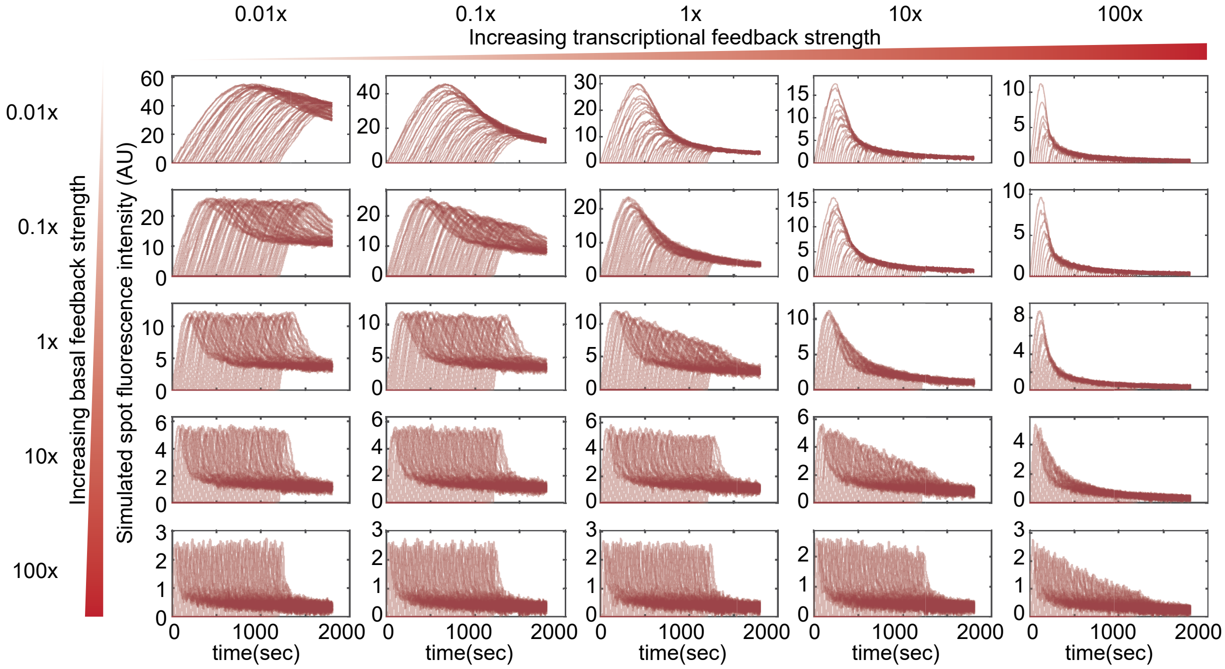

0.01x
0.1x
1x
10x
100x
Increasing transcriptional feedback strength
Increasing basal feedback strength
Simulated spot fluorescence intensity (AU)
time(sec)

**Figure S9: Experiments do not support transcription as the predominant source of negative feedback on NEMO complexes during the primary cytokine response.**

**(A)** Reverse time course experiment testing the effect of transcriptional feedback in trajectories of NEMO complexes. U2OS EGFP-NEMO cells were pretreated for 20 minutes with 50 ng/mL CHX and then stimulated with IL-1 (left) or TNF (right). High-frequency images were collected with the following delay after stimulation: 0 minutes (red), 5 minutes (green), 10 minutes (cyan), or 15 minutes (purple). Only new spots that formed within the first 2 minutes of imaging were tracked to allow direct comparison between early and later-forming EGFP-NEMO complexes. **(B)** Boxplots (median and inter-quartile range) of descriptors of EGFP-NEMO complexes trajectories after stimulation with IL-1 (left) or TNF (right) in cells pretreated with DMSO or CHX. Biological replicates are shown side-by-side with transparency. In nearly all comparisons with early-forming spots (red bin), later forming spots (green, cyan, and purple bins) do not show significant increases ($p$-value >> 0.05, left handed t-test) as predicted by the model where transcriptional negative feedback is predominant.

Fig. S9

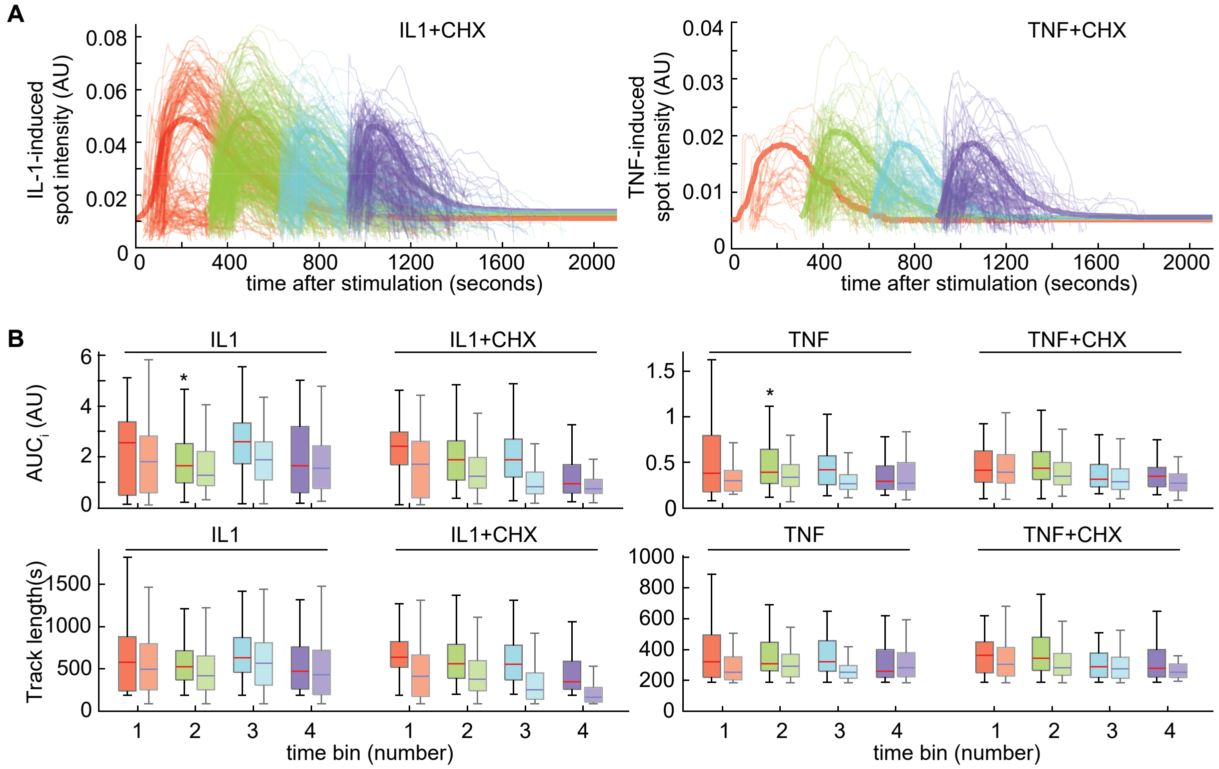
A
IL1+CHX
IL-1-induced spot intensity (AU)
0.08
0.06
0.04
0.02
0
0
400
800
1200
1600
2000
time after stimulation (seconds)
TNF+CHX
TNF-induced spot intensity (AU)
0.04
0.03
0.02
0.01
B
IL1
IL1+CHX
TNF
TNF+CHX
AUC$_i$ (AU)
6
4
2
*
1.5
1
0.5
Track length(s)
1500
1000
500
1000
800
600
400
200
1
2
3
4
time bin (number)

**Figure S10: Validation that CHX concentrations are sufficient to inhibit translation.** **(A)** Cells were pretreated with DMSO (left) or 50 ng/mL CHX (right) for 20 minutes and then stimulated with IL-1. Single-cell time-courses of nuclear RelA do not show appropriate nuclear export in the presence of CHX. **(B)** Bargraphs of $AUC_{fold}$ for trajectories in panel (A) demonstrate that co-stimulation with CHX significantly increases nuclear RelA over the time-course (p-value << 0.05, t-test). These observations are consistent with previous data (Mokashi, Schipper et al. 2019), and together indicate that the NF-κB inhibitory protein IκBα is not being expressed as part of the normal NF-κB transcriptional response.

Fig. S10

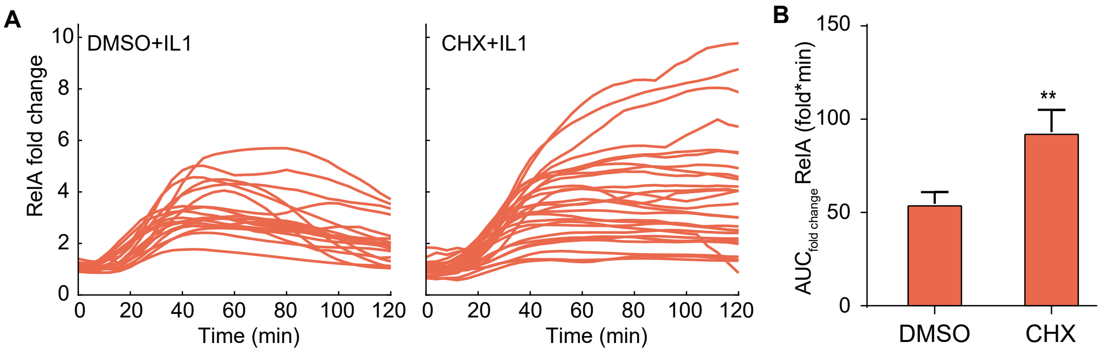
A
DMSO+IL1
CHX+IL1
RelA fold change
Time (min)
B
AUC$_{fold\ change}$ RelA (fold*min)
**
DMSO
CHX

**Figure S11: Hyper-parameter tuning for channel capacity calculations from the variable-gain stochastic pooling network model**

**(A)** Variation of maximum channel capacity as a function of different hyperparameters in the VG-SPN model. Values of the five hyper parameters were tuned to avoid technical biases in the channel capacity calculation. Vertical red lines indicate the value of the hyperparameter used for final model simulations. See supplementary methods for more details. **(B)** Numerical values of channel capacity in bits are presented to accompany Fig. 6E. Although the 'shape' of relationship for different configurations of $CI_{max}$ and gain is consistent between noise models and value of stdR (presented as 'normalized' data in Fig. 6E), the absolute value of the channel capacity in bits is highly dependent on choices for these values. We therefore present bit depth values for reference, but do not contend that the values directly represent capabilities of the biological system.

Fig. S11

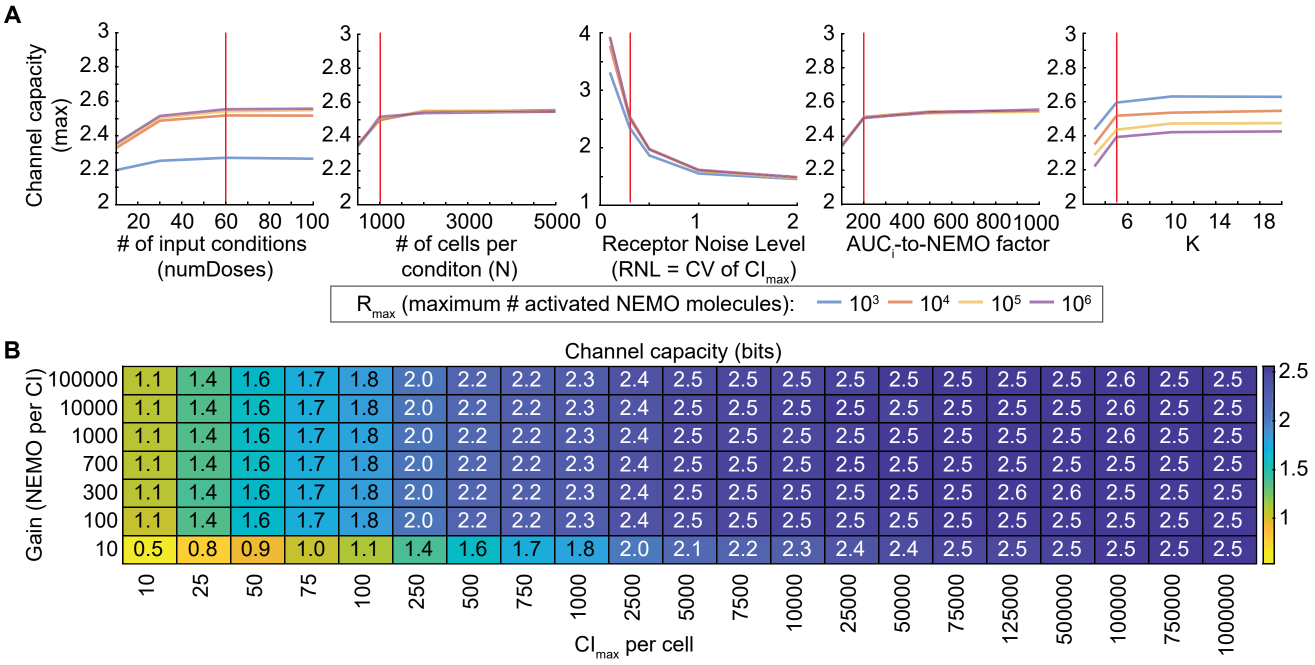
A
Channel capacity (max)
### of input conditions (numDoses)
### of cells per conditon (N)
Receptor Noise Level (RNL = CV of $CI_{max}$)
$AUC_i$-to-NEMO factor
K
$R_{max}$ (maximum # activated NEMO molecules): $10^3$ $10^4$ $10^5$ $10^6$
B
Channel capacity (bits)
Gain (NEMO per CI)
$CI_{max}$ per cell
100000: 1.1 1.4 1.6 1.7 1.8 2.0 2.2 2.2 2.3 2.4 2.5 2.5 2.5 2.5 2.5 2.5 2.5 2.5 2.6 2.5 2.5
10000: 1.1 1.4 1.6 1.7 1.8 2.0 2.2 2.2 2.3 2.4 2.5 2.5 2.5 2.5 2.5 2.5 2.5 2.5 2.6 2.5 2.5
1000: 1.1 1.4 1.6 1.7 1.8 2.0 2.2 2.2 2.3 2.4 2.5 2.5 2.5 2.5 2.5 2.5 2.5 2.5 2.6 2.5 2.5
700: 1.1 1.4 1.6 1.7 1.8 2.0 2.2 2.2 2.3 2.4 2.5 2.5 2.5 2.5 2.5 2.5 2.5 2.5 2.5 2.5 2.5
300: 1.1 1.4 1.6 1.7 1.8 2.0 2.2 2.2 2.3 2.4 2.5 2.5 2.5 2.5 2.5 2.5 2.6 2.6 2.5 2.5 2.5
100: 1.1 1.4 1.6 1.7 1.8 2.0 2.2 2.2 2.3 2.4 2.5 2.5 2.5 2.5 2.5 2.5 2.5 2.5 2.5 2.5 2.5
10: 0.5 0.8 0.9 1.0 1.1 1.4 1.6 1.7 1.8 2.0 2.1 2.2 2.3 2.4 2.4 2.5 2.5 2.5 2.5 2.5 2.5
10 25 50 75 100 250 500 750 1000 2500 5000 7500 10000 25000 50000 75000 125000 500000 100000 750000 1000000

#### Supplementary Movies

**Movie S1. EGFP-NEMO complexes form after IL-1 stimulation, related to Figure 2.**

Colored intensity projection and rotation of 3D time-lapse movie of U2OS EGFP-NEMO cells stimulated with 100 ng/mL IL-1. Each image of the z-stack is colored with a lookup table that goes from blue on the bottom of the cell and becomes progressively redder towards the top of the cell.

**Movie S2. EGFP-NEMO is recruited to punctate complexes after IL-1 and TNF stimulation, related to Figure 2.**

Maximum intensity projection of EGFP-NEMO in cells stimulated with a concentration of 1000 ng/mL for IL-1 (left) and TNF (right).

**Movie S3. Tracking of single EGFP-NEMO complexes after IL-1 stimulation, related to Figure 4.**

Maximum intensity projection of EGFP-NEMO in cells stimulated with 100 ng/mL IL-1. Colored tracks are overlaid for each single-complex trajectory.

#### Supplementary Tables

**Table S1:** Summary of aggregate number of cells and aggregate number of EGFP-NEMO complexed detected, related to Fig. 2 in the main article.

| Stimulation | # cells | # puncta |
|---|---|---|
| **TNF - 0.1 ng/mL** | 33 | 3635 |
| **TNF - 1 ng/mL** | 19 | 20565 |
| **TNF - 10 ng/mL** | 31 | 36213 |
| **TNF - 100 ng/mL** | 32 | 80748 |
| **TNF - 1000 ng/mL** | 14 | 61106 |
| **IL-1 - 0.1 ng/mL** | 37 | 11899 |
| **IL-1 - 1 ng/mL** | 42 | 38389 |
| **IL-1 - 10 ng/mL** | 31 | 87966 |
| **IL-1 - 100 ng/mL** | 35 | 172138 |
| **IL-1 - 1000 ng/mL** | 23 | 152105 |

**Table S2:** Summary of aggregate cell numbers for same cell measurements of EGFP-NEMO and mCh-RELA, related to Fig. 3 in the main article.

| Condition | # cells |
|---|---|
| **Unstimulated** | 28 |
| **TNF - 0.1 ng/mL** | 23 |
| **TNF - 1 ng/mL** | 19 |
| **TNF - 10 ng/mL** | 18 |
| **TNF - 100 ng/mL** | 21 |
| **TNF - 1000 ng/mL** | 22 |
| **IL-1 - 0.1 ng/mL** | 27 |
| **IL-1 - 1 ng/mL** | 26 |
| **IL-1 - 10 ng/mL** | 21 |
| **IL-1 - 100 ng/mL** | 19 |
| **IL-1 - 1000 ng/mL** | 23 |
